# scStable: assessing scRNA-seq analysis stability under bulk-derived between-sample variation

**DOI:** 10.64898/2026.10.01.755876

**Authors:** Chengfeng Jiang, Jingyi Jessica Li

## Abstract

Single-cell RNA sequencing (scRNA-seq) studies often profile only one or a few biological samples, limiting assessment of discovery stability under between-sample variation. We present scStable, a framework for assessing scRNA-seq analysis stability by generating synthetic samples that preserve cell-level heterogeneity from a single-sample reference while injecting bulk-derived between-sample variation. In gastric cancer data comprising 21 matched scRNA-seq and bulk RNA-seq profiles, scStable better recapitulates observed between-sample variation than ZINB-WaVE, subsampling, and Gaussian noise injection. Using synthetic samples, scStable enables stability-aware analysis, including prioritization of stable differentially expressed genes and selection of clustering methods and hyperparameters.

## Background

Single-cell RNA-seq (scRNA-seq) has become a standard technology for characterizing transcriptomes at single-cell resolution [1, 2]. By profiling transcriptomes across thousands to millions of cells, scRNA-seq enables discoveries such as new cell states and their associated marker genes [3, 4]. Despite the growing availability of scRNA-seq data [5], most scRNA-seq studies still profile far fewer biological samples than bulk RNA-seq studies, due to cost constraints [6], limiting the ability to assess whether discoveries generalize to new samples under similar biological conditions. Specifically, a single scRNA-seq sample, regardless of the number of cells profiled, cannot capture between-sample variation, and conclusions derived from one sample may not hold for biologically similar samples [7, 8].

To address the limited number of samples in scRNA-seq studies, method stability is often assessed through resampling-based approaches, such as subsampling cells from a single scRNA-seq sample, including `StabCell` [9] for differentially expressed (DE) gene analysis (i.e., stability to cell sampling), or by varying hyperparameters of a method (i.e., stability to hyperparameter choice)

[10]. However, these approaches cannot reflect stability to between-sample variation, which is essential for generalizable inference [11]. Existing multi-sample scRNA-seq simulators, such as `muscat` [12] and `rescueSim` [13], require multi-sample single-cell data as input and therefore do not apply when only a single scRNA-seq sample is available. As a result, existing scRNA-seq benchmark studies rarely assess method performance stability under realistic between-sample variation [4]. This limitation is consequential: a method may appear highly confident when evaluated on a single sample yet yield substantially different conclusions on biologically similar samples. In practice, analyses often proceed under the implicit assumption that results from one sample are generalizable, even when the magnitude of between-sample variation is unknown [14].

In contrast to scRNA-seq, bulk RNA-seq studies typically have larger sample sizes, partly due to lower cost [15], and have a longer history of statistical modeling for between-sample variation [16, 17]. Bulk RNA-seq samples therefore provide an opportunity to learn between-sample variation that may be used to assess method stability in scRNA-seq data analysis. However, transferring bulk-derived between-sample variation to scRNA-seq data is nontrivial: scRNA-seq captures rich cell-level heterogeneity and exhibits distinct measurement noise and sparsity compared with bulk RNA-seq data [18]. Most existing scRNA-seq simulators that accept a single scRNA-seq sample as input [19–21] focus on mimicking the input sample but are not designed to incorporate bulk-derived between-sample variation [22]. An exception is splatPop [23], which can learn population-level variation from bulk RNA-seq and use it to generate multi-sample scRNA-seq data, but it generates new cells rather than preserving one-to-one correspondence with the cells in the observed scRNA-seq sample (such correspondence is important for cell-level analyses such as cell clustering because it enables tracking the stability of an individual cell’s analysis result). Alternatively, methods such as `bulk2sc` [24] and `scSemiProfiler` [25] infer single-cell expression profiles from bulk RNA-seq samples; however, because bulk RNA-seq aggregates expression across all cells, single-cell profiles are often not uniquely identifiable from bulk data alone, a problem that is especially severe for similar cell types whose highly correlated expression leads to substantial deconvolution uncertainty [26]. The resulting scRNA-seq profiles therefore may not faithfully recover cell-level between-sample variation. Together, existing approaches do not provide synthetic multi-sample scRNA-seq data that both preserve the cells of a given single-sample scRNA-seq reference and incorporate bulk-derived between-sample variation for stability assessment.

Here, we propose scStable, a statistical framework that generates synthetic scRNA-seq samples from a single-sample scRNA-seq reference by injecting bulk-derived between-sample variation while preserving cell-level heterogeneity, enabling stability-aware scRNA-seq analysis when only one scRNA-seq sample is available. The central idea is to leverage a multi-sample bulk RNA-seq reference, either from the same study as the single-sample scRNA-seq reference or, when study-matched bulk data are unavailable, from public repositories such as the Genotype-Tissue Expression (GTEx) project [27], to learn between-sample variation and map it onto the single-sample scRNA-seq reference to generate multiple synthetic scRNA-seq samples. These synthetic samples enable quantification of the stability of discoveries (e.g., DE genes between cell types or biological conditions) under the injected between-sample variation. Moreover, by varying the magnitude of injected between-sample variation, scStable supports stress-testing of computational methods and stability-driven, dataset-specific selection of both methods and their hyperparameters, based on the consistency of analysis results across synthetic scRNA-seq samples. In summary, scStable establishes a bridge between abundant multi-sample bulk RNA-seq data and single-sample scRNA-seq analysis and provides a way to assess the robustness of discoveries from a single sample to between-sample variation.

## Results

### Overview of scStable

scStable is a framework for generating multiple synthetic scRNA-seq samples that preserve the cell-level heterogeneity of a single-sample scRNA-seq reference while injecting bulk-derived between-sample variation learned from a multi-sample bulk RNA-seq reference. These synthetic scRNA-seq samples enable in silico assessment of the stability of discoveries under plausible between-sample variation; they are not substitutes for independently collected biological replicates.

The input to scStable consists of (I) a single-sample scRNA-seq gene-by-cell count matrix (*single-sample scRNA-seq reference*; required) and (II) a multi-sample bulk RNA-seq gene-by-sample count matrix from a matched tissue and condition (*multi-sample bulk RNA-seq reference*; optional). scStable has two modes, which differ in the source of the multi-sample bulk RNA-seq reference. In *bulk-reference mode*, the multi-sample bulk RNA-seq reference comes from the same study as the single-sample scRNA-seq reference, with the multiple samples drawn from the same population. In *bulk-reference-free mode*, no bulk RNA-seq samples are available from the same study, so the multi-sample bulk RNA-seq reference is obtained from tissue-matched data in public repositories such as GTEx [27], either supplied by the user as a count matrix or specified by tissue name, in which case scStable uses parameters estimated from the corresponding GTEx data. The output of scStable is a user-specified number of synthetic scRNA-seq samples. All synthetic scRNA-seq samples contain exactly the same cells as in the single-sample scRNA-seq reference but differ in per-gene expression according to the bulk-derived between-sample variation. Because all synthetic scRNA-seq samples share the same cells, including cell-level covariates, scStable supports cell-level stability assessment (for example, of clustering assignments), unlike cell-resampling approaches and simulators that generate new cells.

The scStable workflow consists of four steps (Fig. 1; full details in Methods). In **Step 1**, scStable fits a statistical model to the single-sample scRNA-seq reference that captures cell-level heterogeneity without requiring cell-type or trajectory annotations, making it possible to assess the stability of cell-level analyses such as clustering and trajectory inference. In **Step 2**, scStable learns how gene expression varies across samples by modeling the multi-sample bulk RNA-seq reference, using study-matched or public data depending on the mode, and estimating bulk-derived between-sample variation. In **Step 3**, scStable transfers this between-sample variation onto the scRNA-seq model to obtain multiple sample-specific models; a user-adjustable scale factor *λ* controls the magnitude of injected variation, with *λ* = 0 corresponding to no injected variation, *λ* = 1 to bulk-derived between-sample variation, and *λ >* 1 to amplified variation for stress-testing. In **Step 4**, scStable generates synthetic scRNA-seq samples from these sample-specific models, keeping the cells fixed across samples while allowing per-gene expression to vary according to the bulk-derived between-sample variation.

**Figure 1:**
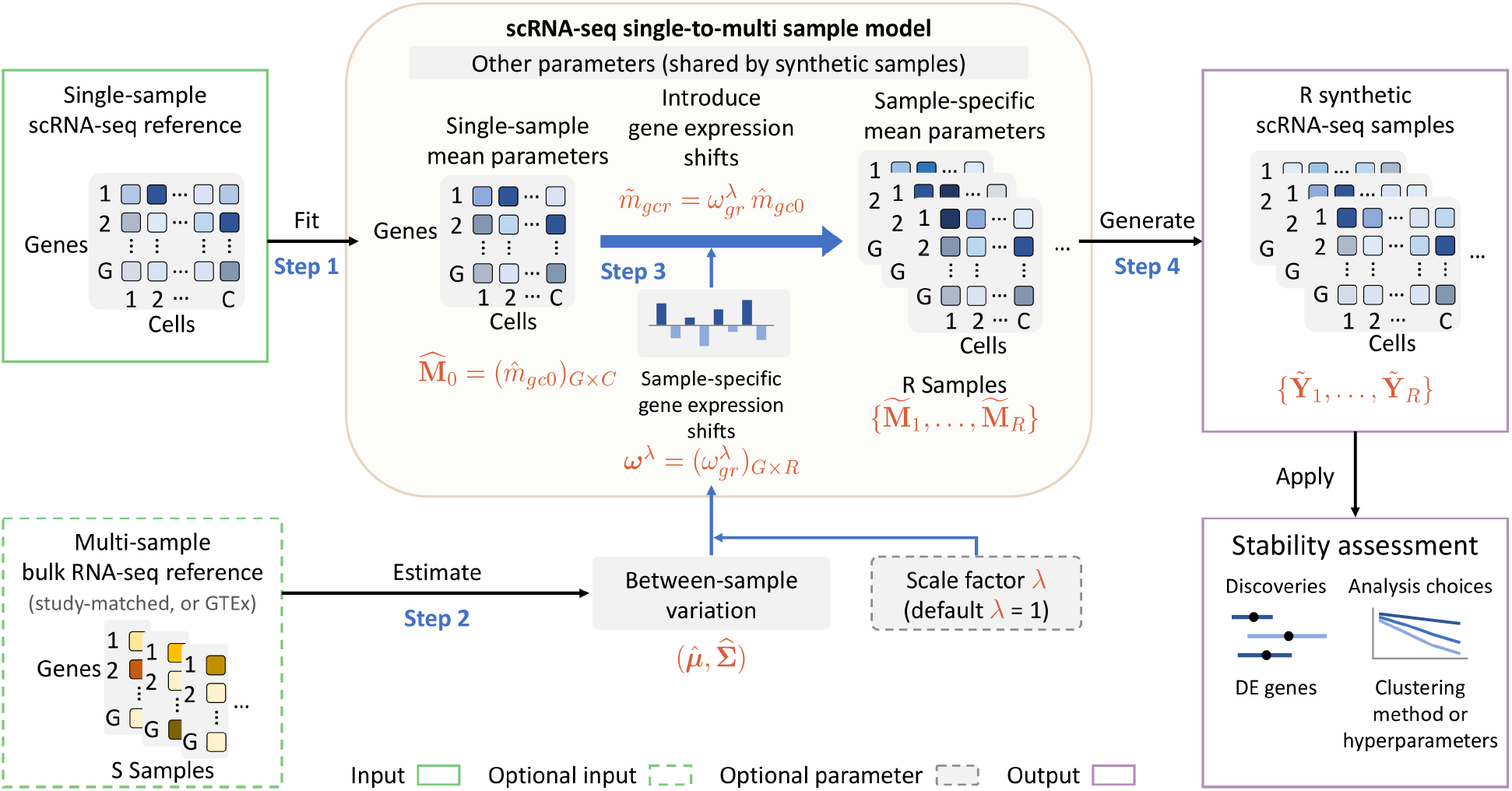
An overview of the scStable workflow. scStable takes as required input a gene-by-cell scRNA-seq count matrix from a *single-sample scRNA-seq reference*. A tissue- and condition-matched *multi-sample bulk RNA-seq reference* can be supplied from the same study as the single-sample scRNA-seq reference (bulk-reference mode); when such a reference is unavailable, scStable can instead use public tissue-matched bulk RNA-seq data such as GTEx, supplied as a count matrix or specified by tissue name (bulk-reference-free mode). **Step 1:** scStable fits a model-based scRNA-seq generative model to the single-sample scRNA-seq reference to estimate the gene-by-cell mean matrix 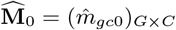 along with additional distributional parameters that capture cell-level heterogeneity and gene–gene dependence (other parameters), which are shared by all synthetic scRNA-seq samples. **Step 2:** scStable models the multi-sample bulk RNA-seq reference to learn bulk-derived between-sample variation, summarized by an estimated mean vector 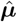 and covariance matrix 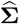 over all *G* genes on the log-CPM scale. **Step 3:** scStable introduces the between-sample variation learned in Step 2 into the single-sample mean parameters from Step 1 through sample-specific gene expression shifts 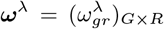, derived from 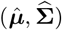, with magnitude controlled by a user-specified scale factor *λ* (default *λ* = 1). For each synthetic scRNA-seq sample, the same gene-specific expression shift is applied across all cells, 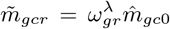, yielding sample-specific mean matrices 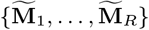 while preserving the cell-level heterogeneity of the reference. **Step 4:** scStable generates *R* synthetic scRNA-seq samples 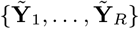 from the resulting sample-specific generative models. Applying the same downstream analysis to each synthetic scRNA-seq sample enables stability-aware assessment of discoveries, such as DE genes, and stability-driven, data-specific selection of analysis methods and hyperparameters. The default *λ* = 1 represents the bulk-derived level of between-sample variation, while varying *λ* enables stress-testing across different variation magnitudes. Because all synthetic scRNA-seq samples share the same set of cells, scStable isolates the effect of between-sample variation while retaining cell-level correspondence across samples.

Because scStable preserves the original cells, cell-type composition is held fixed across synthetic scRNA-seq samples and is not modeled. It is worth noting that the between-sample variation estimated from bulk RNA-seq arises from two sources: expression differences within cell types and differences in cell-type composition across bulk samples. Both sources are represented in the bulk-derived per-gene shifts injected by scStable, so the injected variation can exceed within-cell-type variation alone, making the resulting stability assessment conservative for discoveries intended to generalize across biologically similar samples.

Synthetic scRNA-seq samples generated by scStable support two primary downstream uses. First, they enable *stability-aware assessment* of downstream discoveries by rerunning an analysis pipeline on each synthetic scRNA-seq sample and summarizing how target quantities vary across samples (for example, how a gene’s DE evidence changes across samples). Second, they enable *stability-driven method selection* by comparing candidate methods based on the consistency of their outputs across synthetic scRNA-seq samples, optionally along a path of increasing between-sample variation magnitude for stress testing. Formal definitions of these stability metrics are provided in Methods.

The following Results sections are organized as follows. We first validated that scStable synthetic scRNA-seq samples preserve cell-level heterogeneity while incorporating bulk-derived between-sample variation. We then systematically compared scStable against ZINB-WaVE, which in our implementation preserves one-to-one cell correspondence and has shown strong performance in previous benchmarking studies [28], as well as cell subsampling and Gaussian noise injection. We found that scStable more closely recapitulated real multi-sample scRNA-seq data in terms of between-sample variation at both the pseudo-bulk and single-cell levels. We then demonstrated two applications of scStable: identification of stable DE genes and dataset-specific selection of stable clustering methods and hyperparameters without cell-type labels.

### Validation of scStable synthetic scRNA-seq samples

We first performed a basic validation to assess whether scStable generates synthetic scRNA-seq samples that (i) preserve the cell-level heterogeneity of the single-sample scRNA-seq reference and (ii) reproduce the between-sample variation learned from the multi-sample bulk RNA-seq reference. We used a human peripheral blood mononuclear cell (PBMC) scRNA-seq dataset (commonly referred to as “Zheng4”) [29] as the single-sample scRNA-seq reference, which contains four major immune cell types (Fig. 2A), and GTEx whole-blood bulk RNA-seq with 407 samples as the multi-sample bulk RNA-seq reference. Thus, scStable was run in bulk-reference-free mode. Using scStable, we generated 100 synthetic scRNA-seq samples (details in Supplementary Section S2).

**Figure 2:**
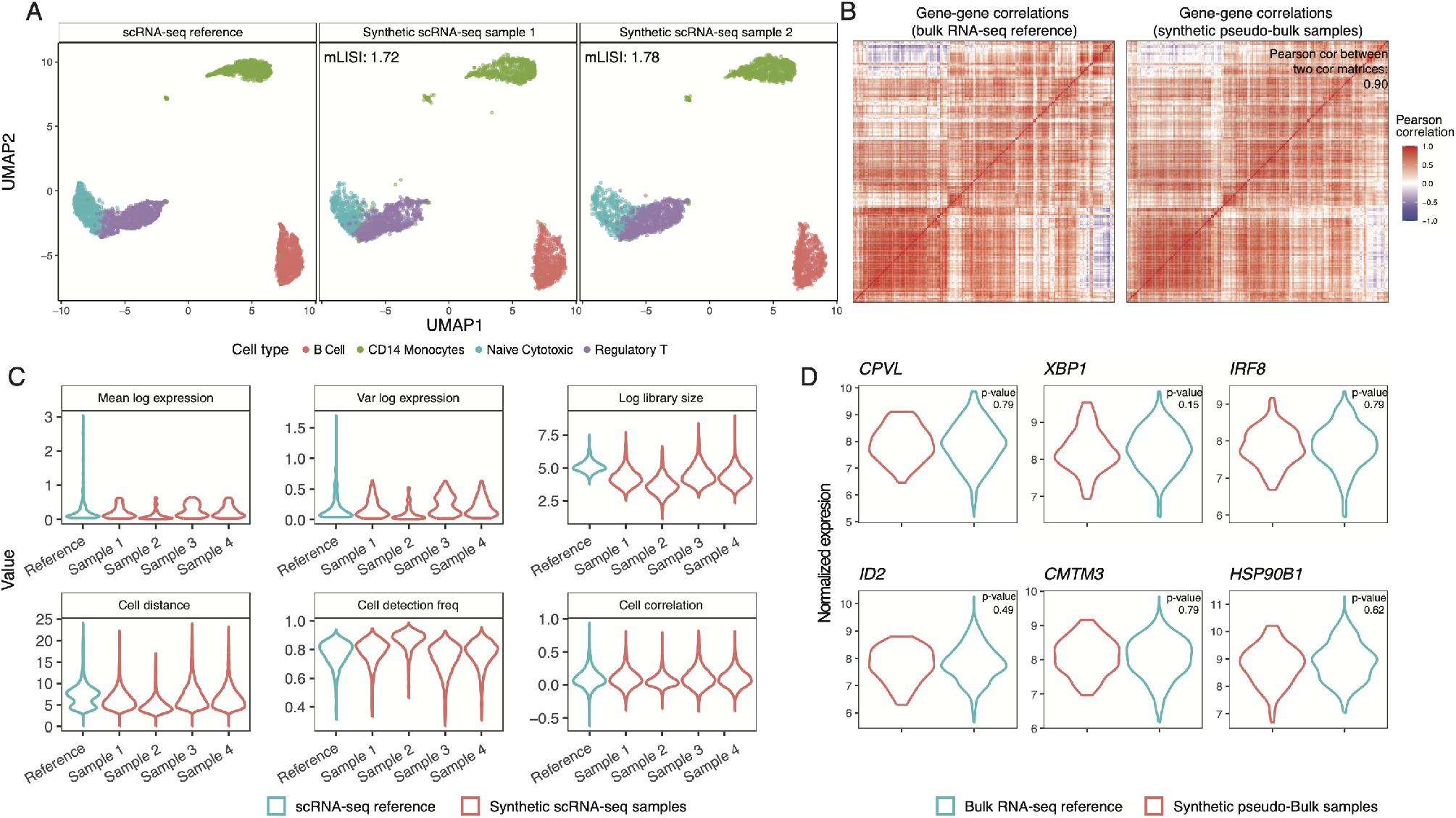
Validation of synthetic scRNA-seq samples generated by scStable. Synthetic scRNA-seq samples preserve the cell-level heterogeneity of the single-sample scRNA-seq reference while incorporating bulk-derived between-sample variation. We use the Zheng4 human PBMC scRNA-seq dataset as the single-sample scRNA-seq reference and GTEx whole-blood bulk RNA-seq as the multi-sample bulk RNA-seq reference. **(A)** UMAP visualizations of the single-sample scRNA-seq reference and two randomly selected synthetic scRNA-seq samples. Four major immune cell types remain well separated, indicating preservation of cell-level heterogeneity. mLISI is the median local inverse Simpson’s index of reference versus synthetic cells in their joint embedding (1.72 and 1.78; maximum 2 when reference and synthetic cells are locally indistinguishable). **(B)** Pearson correlation heatmaps among the top 100 highly variable genes, comparing the multi-sample bulk RNA-seq reference (left) and synthetic pseudo-bulk profiles aggregated from synthetic scRNA-seq samples (right). The correlation structures are highly similar, with a Pearson correlation of 0.90 between the two correlation matrices. **(C)** Violin plots of summary statistics comparing the single-sample scRNA-seq reference (blue) and four randomly selected synthetic scRNA-seq samples (red), including gene-level summaries (mean and variance of log expression) and cell-level summaries (log library size, cell distance, detection frequency, and cell–cell correlation). Summary statistics from synthetic samples show distributions similar to those of the single-sample scRNA-seq reference while exhibiting modest between-sample variation. **(D)** Per-gene marginal distributions on the log-CPM scale for six representative genes, comparing the multi-sample bulk RNA-seq reference (blue; across GTEx samples) and synthetic pseudo-bulk profiles (red; across synthetic scRNA-seq samples). Synthetic pseudo-bulk distributions closely match those of the multi-sample bulk RNA-seq reference for these representative genes, with no significant differences detected by Kolmogorov–Smirnov tests.

To examine preservation of cell-level heterogeneity, we visualized the single-sample scRNA-seq reference and two randomly selected synthetic scRNA-seq samples using uniform manifold approximation and projection (UMAP) in Fig. 2A. The four major immune cell types remained similarly well separated in the synthetic scRNA-seq samples as in the reference, indicating that scStable preserves the cell-level heterogeneity of the scRNA-seq data, with no evidence of spurious splitting or unintended mixing across cell types. This observation was further supported by high median local inverse Simpson’s index (mLISI) values of 1.72 and 1.78 for the two synthetic scRNA-seq samples when each was jointly analyzed with the single-sample scRNA-seq reference (the maximum possible value is 2), indicating strong similarity to the reference.

Looking more closely, summary statistics computed on four randomly selected synthetic scRNA-seq samples showed distributions similar to those of the single-sample scRNA-seq reference, including cell-level statistics (e.g., log library size, cell distance, detection frequency, and cell–cell correlation) and gene-level statistics (e.g., mean and variance of log expression) (Fig. 2C). At the same time, modest differences in these distributions were observed across synthetic scRNA-seq samples and relative to the single-sample reference, as expected from the injected between-sample variation.

Because Zheng4 contains only a single scRNA-seq sample, no real multi-sample scRNA-seq data are available for direct validation of the injected between-sample variation. We therefore performed a sanity check at the pseudo-bulk level against the same multi-sample bulk RNA-seq reference from which scStable learned the between-sample variation. For each synthetic scRNA-seq sample, we constructed a pseudo-bulk profile by aggregating counts across cells and aligned the pseudo-bulk to the multi-sample bulk RNA-seq reference on the logarithmic counts per million (log-CPM) scale, as described in Step 3 of Methods. For illustration, Fig. 2D shows six representative genes for which the marginal distributions of synthetic pseudo-bulk profiles had similar means and variances to those observed in the multi-sample bulk RNA-seq reference, with no significant differences detected by Kolmogorov–Smirnov (KS) tests. Beyond per-gene marginal distributions, the gene–gene correlations observed in the multi-sample bulk RNA-seq reference were well preserved in the synthetic pseudo-bulk samples (Fig. 2B), with a Pearson correlation of 0.90 between the two sets of correlation values. Hence, this analysis provides a sanity check that scStable preserves cell-level heterogeneity and transfers the intended bulk-derived between-sample variation to the synthetic scRNA-seq samples. A more direct validation against real multi-sample scRNA-seq data is presented in the next section.

### scStable better captures between-sample variation than ZINB-WaVE, subsampling, and Gaussian noise injection

We next performed a more direct validation of whether scStable recapitulates the between-sample variation observed in real multi-sample scRNA-seq data. We used a paired gastric cancer dataset [30] containing matched bulk and scRNA-seq profiles from the same individuals. After filtering to 21 individuals with both data types (Methods), we designated one scRNA-seq sample as the single-sample scRNA-seq reference and retained the remaining 20 scRNA-seq samples as a held-out empirical reference for real between-sample variation. All 21 paired bulk RNA-seq profiles were used as the multi-sample bulk RNA-seq reference from which scStable learned between-sample variation. This design allowed us to evaluate whether variation learned from bulk RNA-seq and transferred by scStable reproduced the variation actually observed across real scRNA-seq samples.

For scStable, we considered two settings of its variation scale parameter *λ*: *λ* = 1, corresponding to the bulk-derived between-sample variation, and *λ* = 2, representing amplified variation for stress testing. We compared both settings with ZINB-WaVE, Gaussian noise injection, and cell-wise subsampling at both the pseudo-bulk and single-cell levels (Table 1; Methods). ZINB-WaVE was selected as the simulator comparator because it preserves gene identities and, in our implementation, one-to-one cell correspondence with the reference, enabling direct per-gene and per-cell comparisons. The Gaussian noise injection approach generated synthetic samples by adding independently drawn gene-specific Gaussian shifts on the log-expression scale to all cells of the reference, with noise magnitudes estimated from the observed cell-to-cell variation of each gene. Cell-wise subsampling generated each synthetic sample by randomly retaining 80% of the reference cells without replacement, thereby capturing cell-sampling variability but not biological between-sample variation.

**Table 1:**
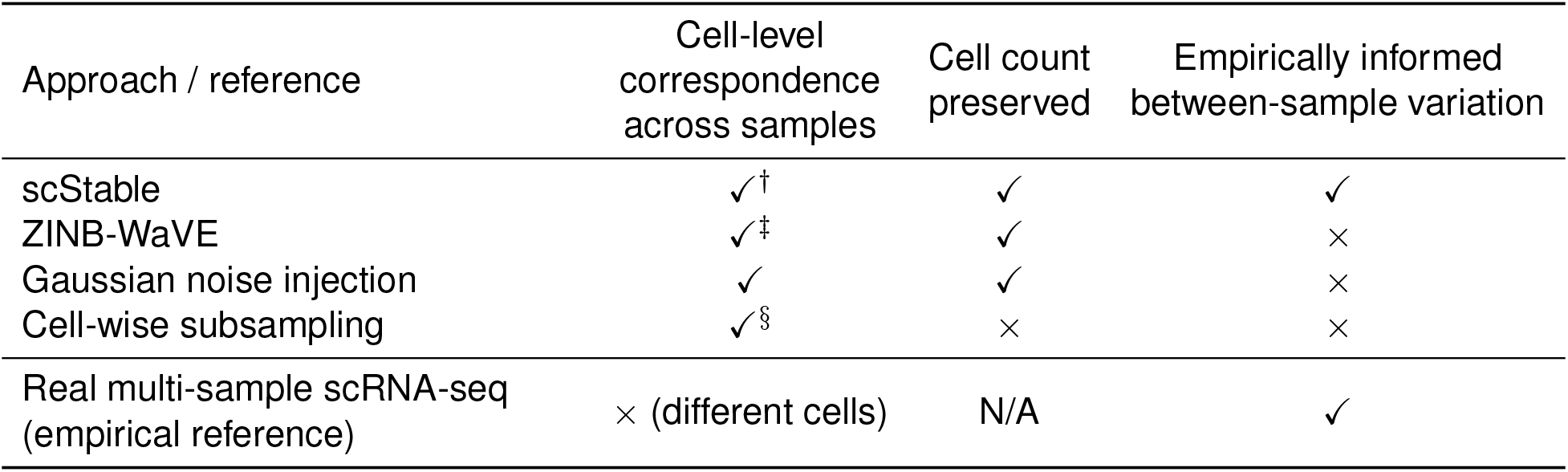
Comparison of scStable and alternative approaches used in this study for generating multiple synthetic scRNA-seq samples from a single-sample scRNA-seq reference. Real multi-sample scRNA-seq from the paired gastric cancer dataset [30] is included as the empirical reference and is not a generative approach. *Cell-level correspondence across samples*: whether the generated samples retain one-to-one correspondence with the cells in the single-sample scRNA-seq reference, enabling per-cell stability analysis. *Cell count preserved* : whether each generated sample retains the full set of cells from the single-sample scRNA-seq reference. *Empirically informed between-sample variation*: whether differences across samples reflect empirically observed between-sample variation rather than variation arising solely from model stochasticity, resampling, or arbitrary noise injection. ^*†*^scStable retains the reference cells and their PCA-derived cell-level covariates across synthetic samples and does not require cell-type labels as input. ^*‡*^In our implementation, ZINB-WaVE synthetic scRNA-seq samples are generated from the fitted model while retaining the fitted cell-specific parameters, so that each simulated cell corresponds one-to-one to a cell in the single-sample scRNA-seq reference (Methods). ^*§*^Cell-wise subsampling retains only a subset of the reference cells in each synthetic scRNA-seq sample, so cell-level correspondence is only partial.

All pseudo-bulk-level statistics below were computed after normalizing library size, so that differences in cell number across real samples were not counted as between-sample variation (Methods; the performance differences between scStable and the alternatives remained similar without this step, Supplementary Fig. S2). At the pseudo-bulk level, increasing *λ* from 1 to 2 had little effect on the concordance of per-gene across-sample means with those of the held-out real multi-sample scRNA-seq data (Lin’s concordance *ρ*_*c*_ = 0.902 and 0.891, respectively), but improved the concordance of per-gene between-sample variances with the real data (from *ρ*_*c*_ = 0.774 to 0.874; Fig. 3A). The total between-sample variance relative to real multi-sample scRNA-seq also increased from 0.45 at *λ* = 1 to 0.98 at *λ* = 2 (Supplementary Fig. S1). Thus, as expected, *λ* controls the magnitude of between-sample variation generated by scStable without substantially altering the per-gene mean structure.

**Figure 3:**
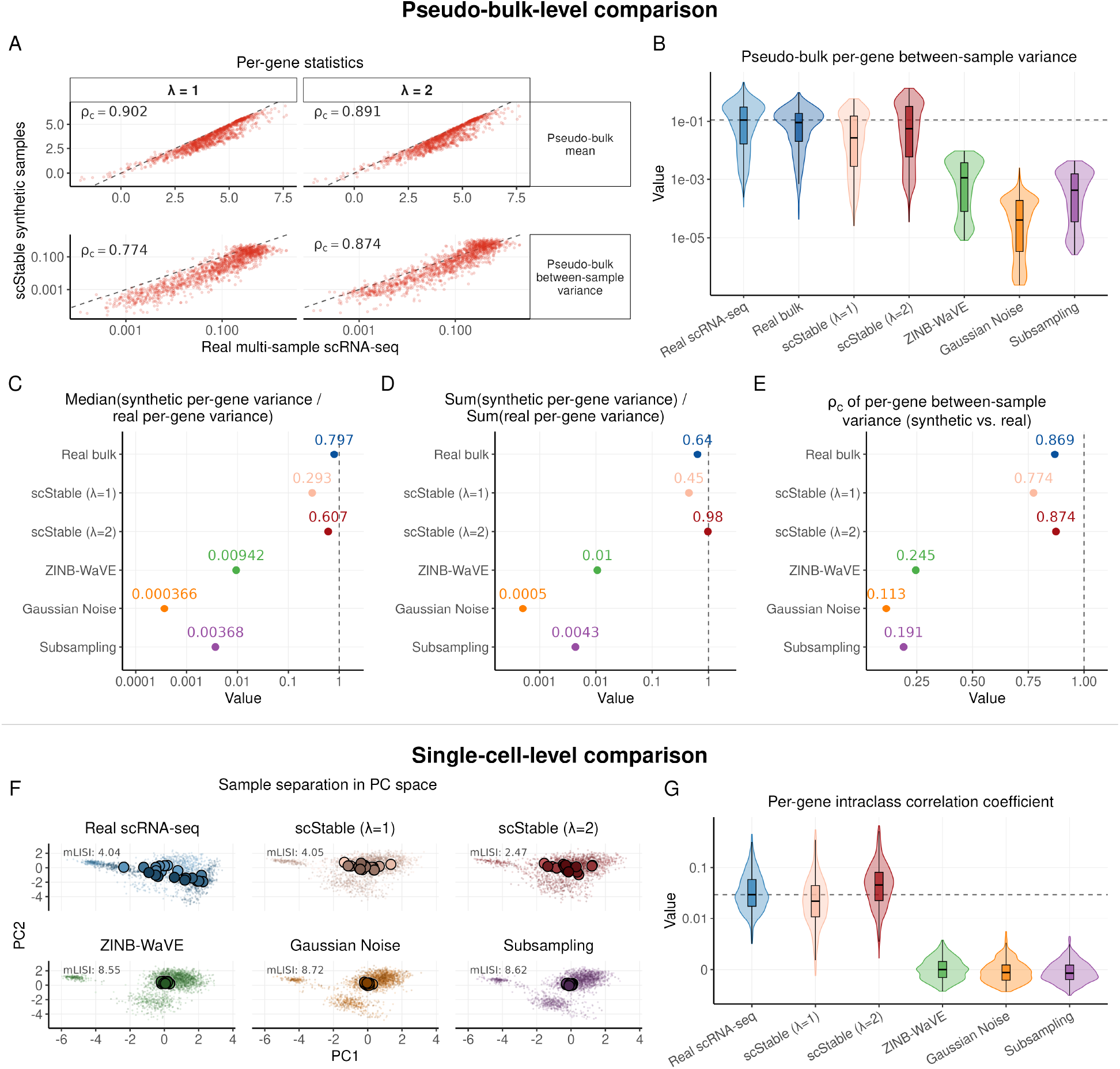
scStable better captures between-sample variation than ZINB-WaVE, subsampling, and Gaussian noise injection. Using a paired gastric cancer dataset with multi-sample scRNA-seq and bulk RNA-seq from the same individuals, we assess how well scStable reproduces the between-sample variation observed in held-out real multi-sample scRNA-seq and compare it with ZINB-WaVE, Gaussian noise injection, and cell-wise subsampling. Between-sample variation is evaluated at the pseudo-bulk level (A–E) and single-cell level (F–G). **(A)** Per-gene pseudo-bulk mean (top) and between-sample variance (bottom; log_10_ scale) of scStable synthetic scRNA-seq samples at *λ* = 1 and *λ* = 2, compared with real multi-sample scRNA-seq. Each point represents one gene; the dashed line is *y* = *x*, and *ρ*_*c*_ denotes Lin’s concordance correlation coefficient. **(B)** Distributions of per-gene between-sample variance for real multi-sample scRNA-seq, paired bulk RNA-seq, and synthetic scRNA-seq samples generated by each approach. The dashed line indicates the median for real multi-sample scRNA-seq. **(C)** Median across genes of the ratio of each approach’s per-gene between-sample variance to that of real multi-sample scRNA-seq. **(D)** Total between-sample variance relative to real multi-sample scRNA-seq. In (C,D), the dashed line at 1 indicates exact agreement. **(E)** *ρ*_*c*_ between per-gene between-sample variances from each approach and those from real multi-sample scRNA-seq; the dashed line at 1 indicates exact agreement. **(F)** Cells from each approach projected into a common principal-component (PC) space. Small points represent individual cells and large outlined points represent sample centroids in the PC space. For comparability, 20 samples are used for each approach. mLISI is calculated over sample identity and ranges from 1, when samples are locally separated, to 20, when samples are locally indistinguishable. **(G)** Per-gene intraclass correlation coefficients, measuring the fraction of cell-level expression variance attributable to sample identity. The dashed line indicates the median for real multi-sample scRNA-seq.

We next compared how well the generated samples from different approaches recapitulated the between-sample variation observed in the held-out real scRNA-seq samples. At both *λ* = 1 and 2, the distribution of per-gene between-sample variances generated by scStable was similar to that of the paired bulk RNA-seq samples, as expected because scStable learns between-sample variation from these samples. Importantly, the scStable distributions were also substantially closer to the distribution observed in the held-out real multi-sample scRNA-seq samples than those generated by ZINB-WaVE, Gaussian noise injection, or cell-wise subsampling (Fig. 3B). For each gene, we computed the ratio of the between-sample variance generated by each approach to that observed in real multi-sample scRNA-seq and summarized these ratios by their median across genes, with values closer to 1 indicating better agreement. The median ratio was 0.607 for scStable at *λ* = 2 and 0.293 at *λ* = 1, compared with 0.00942 for ZINB-WaVE, 0.00368 for subsampling, and 0.000366 for Gaussian noise injection; paired bulk RNA-seq itself reached 0.797 (Fig. 3C). In aggregate, scStable recovered 0.98 of the total between-sample variation across genes in real multi-sample scRNA-seq at *λ* = 2 and 0.45 at *λ* = 1, compared with 0.01, 0.0043, and 0.0005 for ZINB-WaVE, subsampling, and Gaussian noise injection, respectively, while paired bulk RNA-seq reached 0.64 (Fig. 3D). Agreement of per-gene between-sample variances with the real scRNA-seq samples showed a similar pattern: Lin’s concordance was *ρ*_*c*_ = 0.874 for scStable at *λ* = 2 and 0.774 at *λ* = 1, close to that of paired bulk RNA-seq (0.869), compared with 0.245 for ZINB-WaVE, 0.191 for subsampling, and 0.113 for Gaussian noise injection (Fig. 3E).

The availability of held-out real scRNA-seq samples also allowed us to evaluate between-sample variation at the single-cell level, which was not possible in the Zheng4 analysis above. We embedded all six sets of scRNA-seq samples—real multi-sample scRNA-seq, scStable at *λ* = 1, scStable at *λ* = 2, ZINB-WaVE, Gaussian noise injection, and cell-wise subsampling—in a common principal-component space constructed from their pooled cells. The sample centroids of real multi-sample scRNA-seq were dispersed across the space, whereas those from ZINB-WaVE, Gaussian noise injection, and cell-wise subsampling collapsed toward essentially a single point, indicating substantial underrepresentation of between-sample variation and consistent with the pseudo-bulk-level results (Fig. 3F). In contrast, scStable reproduced substantially greater sample-to-sample separation, with greater separation at *λ* = 2 than at *λ* = 1. We quantified sample mixing using the median local inverse Simpson’s index (mLISI; Methods) over sample labels, which measures the effective number of samples represented among a cell’s neighbors and approaches 1 when samples are separable and 20 when samples are interchangeable. Real multi-sample scRNA-seq had an mLISI of 4.04, while ZINB-WaVE, Gaussian noise injection, and cell-wise subsampling had values of 8.55, 8.72, and 8.62, respectively; scStable reached 4.05 at *λ* = 1 and 2.47 at *λ* = 2, substantially closer to the real multi-sample scRNA-seq data than the other generative approaches. Decomposition of cell-level variance using the intraclass correlation coefficient (ICC; Methods) showed a consistent pattern: the median ICC across genes was 3.0% in the real multi-sample scRNA-seq data, compared with 2.2% and 4.6% under scStable at *λ* = 1 and *λ* = 2, respectively, whereas the corresponding values for the other approaches were near zero (|ICC| *<* 0.05%; Fig. 3G).

Together, these results provide a direct validation of scStable against held-out real multi-sample scRNA-seq data. Among the approaches compared, scStable most closely reproduced between-sample variation in the real data at both the pseudo-bulk and single-cell levels, across both per-gene and aggregate summaries. The two levels responded differently to *λ*: *λ* = 2 closely matched the total pseudo-bulk variance of real multi-sample scRNA-seq, whereas *λ* = 1 more closely matched the real data at the single-cell level, suggesting that the unamplified bulk-derived variation provides a reasonable default for this dataset. This discrepancy likely reflects a limitation of transferring bulk-derived variation to a fixed set of cells: scStable preserves the cell composition of the single-sample scRNA-seq reference, whereas real between-sample variation includes both within-cell-type expression changes and changes in cell-type composition. Because bulk RNA-seq does not directly resolve these two sources of variation, exact agreement with real multi-sample scRNA-seq at both the pseudo-bulk and single-cell levels is not expected.

### Application 1: scStable enables identification of stable DE genes

We demonstrate how scStable enables assessment of the stability of DE genes under between-sample variation. In both between-cell-type and between-condition DE analyses, we used synthetic scRNA-seq samples generated by scStable to quantify how stable DE evidence is under bulk-derived between-sample variation. For each gene, we computed a quantile interval of its DE evidence, summarized by the Benjamini–Hochberg (BH)-adjusted *p*-value, across synthetic scRNA-seq samples, with smaller values indicating stronger evidence for differential expression. We then identified DE genes that were significant in the reference scRNA-seq data and stable across synthetic scRNA-seq samples.

We first considered between-cell-type DE using the Zheng4 dataset, focusing on regulatory T cells versus B cells. We applied six commonly used scRNA-seq DE methods implemented in Seurat v5 [31] to the single-sample scRNA-seq reference and to each synthetic scRNA-seq sample generated by scStable, using the same GTEx whole-blood multi-sample bulk RNA-seq reference as in the validation analysis. For each DE method and each gene, we summarized stability by the length of the adjusted *p*-value quantile interval across synthetic scRNA-seq samples (Methods), with shorter intervals indicating greater stability. Fig. 4A (bottom) shows the distribution of interval lengths among DE genes that were significant in the single-sample scRNA-seq reference (adjusted *p*-value *<* 0.05). While most DE genes had short intervals, a subset showed markedly wider intervals, indicating that their DE evidence was sensitive to between-sample variation despite being significant in the reference. DE genes with interval length below 0.05 were classified as *stable DE genes*, while the remaining significant DE genes were referred to as *other DE genes* (Methods). Fig. 4A (top) illustrates this contrast using representative genes. We next examined the relationship between DE strength in the reference sample and stability across synthetic samples. For each DE method, we ranked genes by adjusted *p*-value in the reference sample from smallest to largest (*DE Rank* ) and separately by quantile-interval length from smallest to largest (*Stability Rank* ). Fig. 4B shows that although many genes ranked highly under both criteria, some genes with strong DE evidence in the reference sample dropped substantially in Stability Rank, indicating that their DE evidence was less stable under between-sample variation. For example, under the Wilcoxon rank-sum test, *GAPDH* changed from DE Rank 134 to Stability Rank 271, while *SERF2* changed from 128 to 427 and *SH3BGRL3* from 143 to 429. In contrast, genes ranked highly under both criteria represented strong DE signals in the reference whose DE evidence was also relatively stable across synthetic samples. KEGG enrichment analysis further showed that stable DE genes were enriched for immune-related pathways consistent with the regulatory T-cell versus B-cell comparison, whereas other DE genes showed different and generally less cell-type-specific enrichment patterns (Fig. 4C).

**Figure 4:**
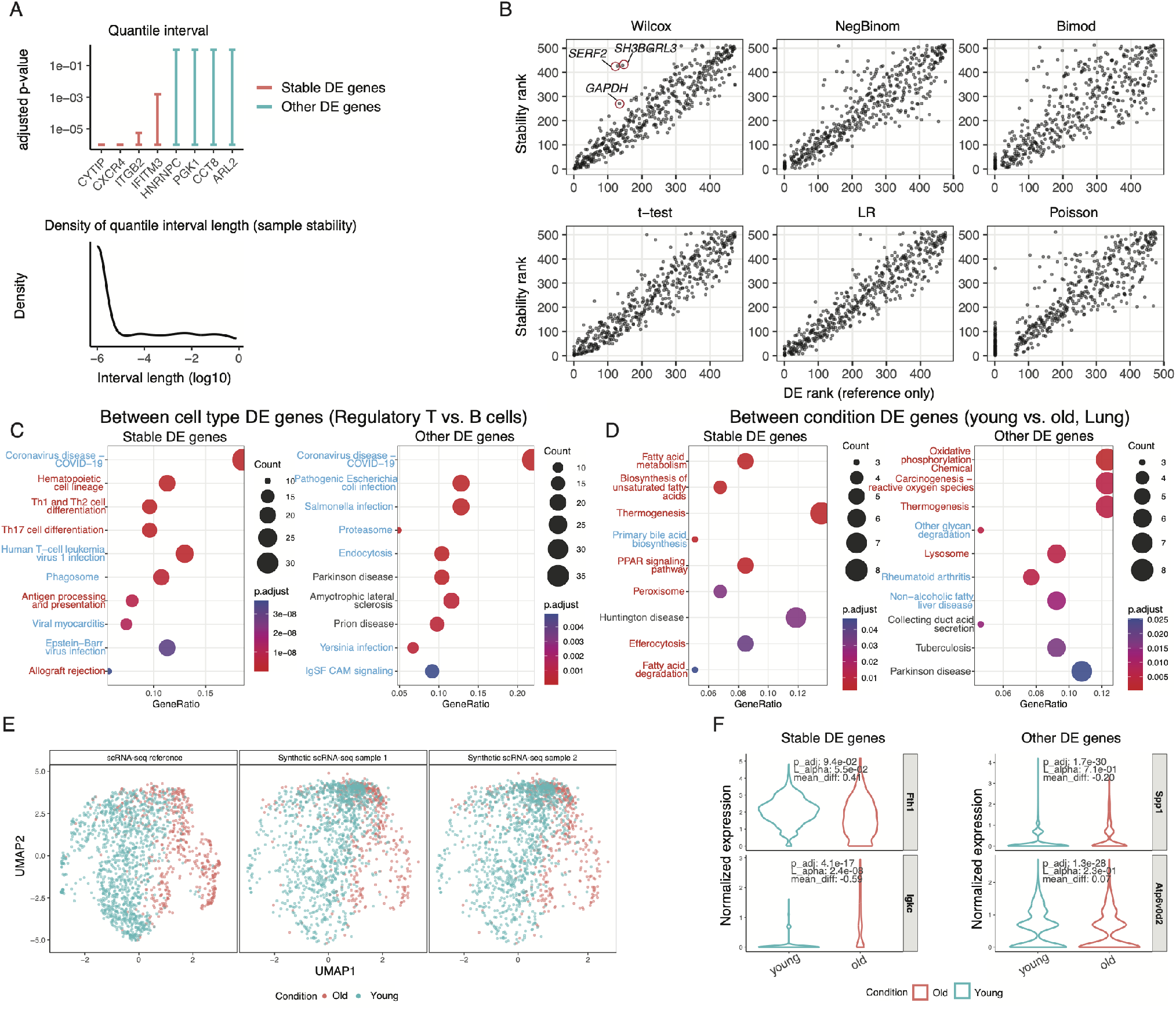
Identification and prioritization of stable differentially expressed (DE) genes using scStable synthetic scRNA-seq samples. Stability analysis is shown for DE between cell types (Zheng4 human PBMC) and between conditions (lung aging). **(A)** Top: 95% quantile intervals of per-gene BH-adjusted *p*-values across synthetic scRNA-seq samples. Among genes significant in the single-sample scRNA-seq reference (*p*_adj_ *<* 0.05), genes with quantile interval length *L*_*α*_ *<* 0.05 are defined as stable DE genes (red), whereas other DE genes are shown in teal. Bottom: distribution of *L*_*α*_ among genes significant in the single-sample scRNA-seq reference. **(B)** Between-cell-type DE (Regulatory T vs. B cells, Zheng4). Gene ranks based on DE evidence in the single-sample scRNA-seq reference (x-axis; *DE Rank* ) are compared with ranks based on stability across synthetic scRNA-seq samples (y-axis; *Stability Rank* ) for six scRNA-seq DE methods. Red-circled genes (GAPDH, SERF2, SH3BGRL3) rank substantially lower by stability than by DE evidence. **(C**,**D)** KEGG pathway enrichment comparing stable and other DE genes for between-cell-type (C) and between-condition (D) analyses. **(E)** Between-condition DE (young vs. old, lung). UMAPs of the pooled age-specific reference scRNA-seq data and two pairs of synthetic scRNA-seq samples representing the young and old conditions show preservation of cell-level heterogeneity and condition separation. **(F)** Violin plots of normalized expression for representative stable and other DE genes. Stable DE genes show more consistent between-condition differences across synthetic scRNA-seq samples. For each gene, the reference adjusted *p*-value (p adj), quantile interval length (*L*_*α*_), and difference in mean expression between young and old conditions (mean diff) are reported.

We next demonstrated condition-level stability analysis using macrophages from an aging mouse lung dataset [32]. Because individual mice contributed too few macrophage cells for stable model fitting, macrophage cells were pooled across mice within each age group to construct age-specific scRNA-seq references for young and old mice, respectively (Methods). This analysis therefore serves as a proof-of-concept demonstration of condition-level stability assessment within a single cell type rather than a literal single-biological-sample application. We ran scStable separately using the young and old pooled scRNA-seq references, with the corresponding multi-sample flow-sorted macrophage bulk RNA-seq data used to learn between-sample variation for each age group. This generated two sets of synthetic scRNA-seq samples representing the young and old conditions.

Fig. 4E shows that the synthetic scRNA-seq samples preserved cell-level heterogeneity while maintaining the global separation between the young and old age-specific pooled reference data. We performed cell-level DE analysis between the young and old pooled references using the Wilcoxon rank-sum test and repeated the same comparison across the corresponding synthetic samples. As above, for each gene we constructed a quantile interval of adjusted *p*-values across synthetic comparisons and classified genes that were significant in the pooled reference comparison (adjusted *p*-value *<* 0.05) as stable DE genes if their interval length was below 0.05, and as other DE genes otherwise. Because cells were pooled across mice within each age group, this analysis is intended to demonstrate the stability-assessment workflow rather than provide population-level inference on age effects across animals.

Representative examples in Fig. 4F show that stable DE genes exhibited more consistent expression differences between young and old conditions across synthetic samples, whereas other DE genes showed less stable differences under the modeled between-sample variation. Consistent with this distinction, KEGG enrichment analysis showed different enrichment patterns between stable and other DE genes (Fig. 4D), with stable DE genes highlighting pathways plausibly related to aging-associated biological differences. Together, these analyses demonstrate how scStable enables stability-aware DE analysis by distinguishing DE signals that remain robust under bulk-derived between-sample variation from those that are more sensitive to such variation.

### Application 2: scStable enables data-specific selection of stable clustering methods without requiring cell-type labels

We used scStable synthetic scRNA-seq samples to study cell clustering stability under bulk-derived between-sample variation and to select clustering methods and their hyperparameters based on stability across synthetic scRNA-seq samples, without requiring known cell-type labels. Using the human PBMC Zheng4 dataset as the single-sample scRNA-seq reference and GTEx whole-blood bulk RNA-seq as the multi-sample bulk RNA-seq reference, we generated synthetic scRNA-seq samples under a range of between-sample variation magnitudes controlled by the scale factor *λ* (Methods, Step 3). Smaller values of *λ* produce synthetic scRNA-seq samples closer to the single-sample scRNA-seq reference, whereas larger values induce stronger between-sample variation, enabling systematic stress-testing of clustering stability. Compared with cell subsampling approaches, scStable introduces between-sample variation learned from the multi-sample bulk RNA-seq reference. Moreover, because all synthetic scRNA-seq samples contain the same cells with shared cell-level covariates, scStable isolates the effect of between-sample variation and enables cell-level stability assessment, which is not possible with cell subsampling approaches. We considered three aspects of clustering stability: identifying cells with unstable cluster assignments, comparing clustering methods by stability, and selecting hyperparameters for the most stable method.

We first identified unstable cell assignments using an entropy-based per-cell stability score derived from the consensus co-clustering matrix across synthetic scRNA-seq samples at *λ* = 1, corresponding to bulk-derived between-sample variation (Methods), with higher scores indicating more decisive and consistent cluster assignments. Fig. 5A shows that most cells exhibited high stability, indicating consistent cluster assignments under between-sample variation. In contrast, cells from the naive cytotoxic and regulatory T populations showed noticeably lower stability, particularly among cells that were difficult to distinguish as naive cytotoxic versus regulatory T cells based on their transcriptional profiles, indicating that their cluster assignments were more sensitive to between-sample variation. By comparison, B cells and CD14 monocytes, which formed well-separated clusters, exhibited uniformly high stability scores. These results illustrate how scStable synthetic scRNA-seq samples can identify cells with unstable cluster assignments that may warrant cautious interpretation during cell-type annotation.

**Figure 5:**
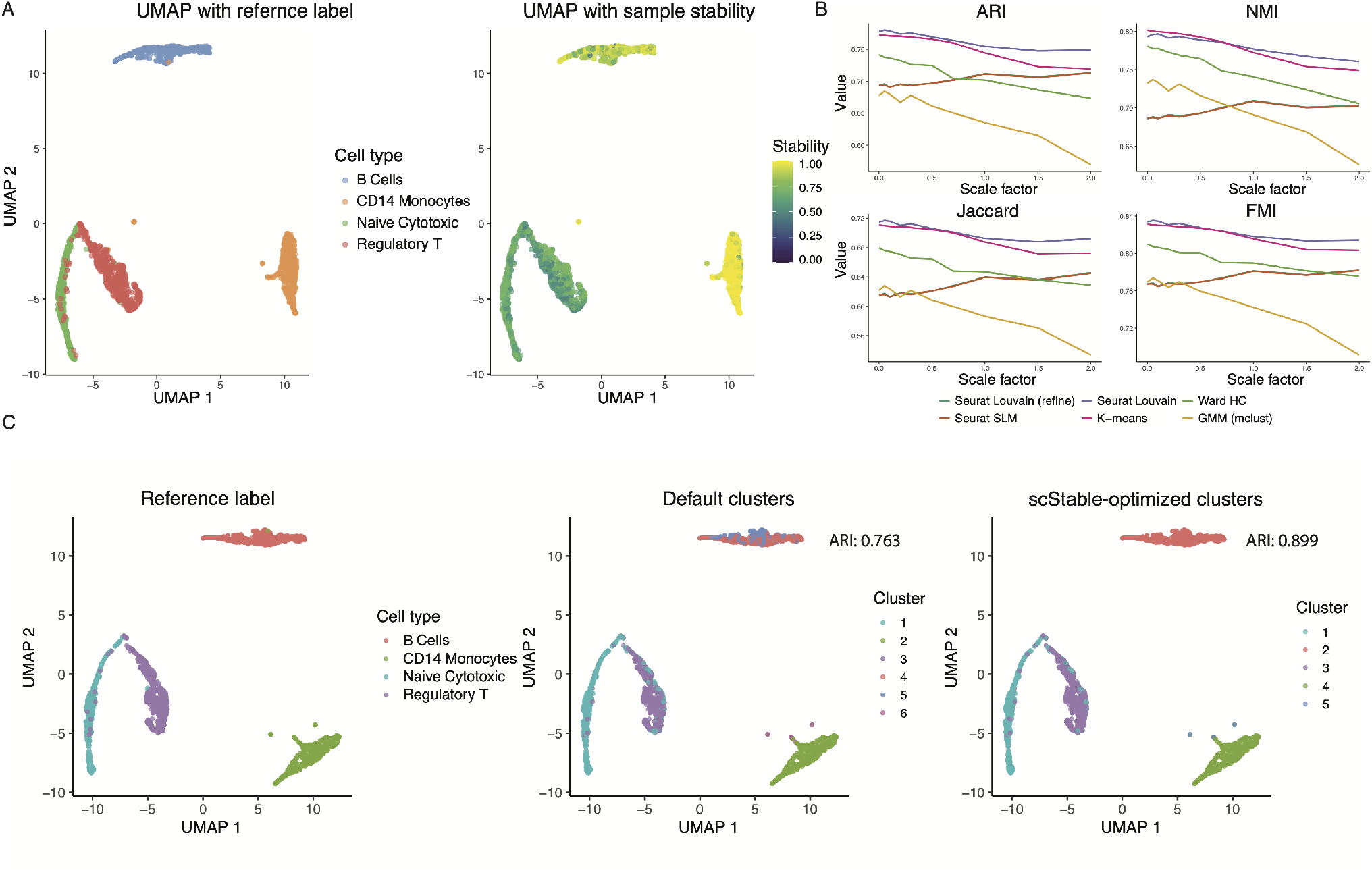
Data-specific clustering stability analysis using scStable synthetic scRNA-seq samples. scStable synthetic scRNA-seq samples are used to identify cells with unstable cluster assignments under bulk-derived between-sample variation and to select clustering methods and hyperparameters based on their stability, without requiring known cell-type labels. The Zheng4 human PBMC scRNA-seq dataset is used as the single-sample scRNA-seq reference, with GTEx whole-blood bulk RNA-seq as the multi-sample bulk RNA-seq reference. **(A)** UMAP visualization of the single-sample scRNA-seq reference colored by known cell-type labels (left) and by entropy-based clustering stability (right), computed from the consensus matrix across synthetic scRNA-seq samples at *λ* = 1. Cells with lower stability tend to occur in regions where the naive cytotoxic and regulatory T populations are transcriptionally less well separated. Known cell-type labels are shown only for evaluation and are not used to compute the stability scores. **(B)** Stability of clustering methods under increasing magnitudes of between-sample variation, controlled by the scale factor *λ*. Methods evaluated include Louvain, Louvain refinement, and Smart Local Moving (SLM) from Seurat; K-means and Ward hierarchical clustering (Ward HC); and Gaussian mixture modeling (GMM) from mclust. For each method and each *λ*, clustering labels from synthetic scRNA-seq samples are compared with those from the single-sample scRNA-seq reference using Adjusted Rand Index (ARI), Normalized Mutual Information (NMI), Jaccard Index, and Fowlkes–Mallows Index (FMI). Methods whose agreement remains high as *λ* increases are more stable for this dataset. **(C)** UMAP visualization comparing known cell-type labels with Louvain clustering results obtained using the default hyperparameters and the hyperparameter combination selected based on stability across scStable synthetic scRNA-seq samples (*n*_PC_ = 10, *k*_NN_ = 20, *r* = 0.40; *λ* = 1). Compared with the default hyperparameters, the stability-selected hyperparameters yield clustering with greater correspondence to the known cell-type labels, with ARI increasing from 0.763 to 0.899. Known cell-type labels are used only for evaluation and not for hyperparameter selection.

We next examined how scStable enables data-specific selection of clustering methods by comparing their stability under increasing between-sample variation. Without requiring cell-type labels, we assessed stability by comparing clustering results from each synthetic scRNA-seq sample with those from the single-sample scRNA-seq reference, thereby measuring the sensitivity of each method’s clustering result to modeled between-sample variation. We evaluated six commonly used clustering methods: Louvain, Louvain refinement, and Smart Local Moving (SLM) from Seurat v5; K-means and Ward hierarchical clustering in base R; and Gaussian mixture modeling (GMM) from the `mclust` package [33]. For each method and each value of *λ*, we clustered each synthetic scRNA-seq sample and measured agreement with the reference clustering using adjusted Rand index (ARI), normalized mutual information (NMI), Jaccard index, and Fowlkes– Mallows index (FMI) (Fig. 5B). Across all four metrics, Louvain clustering maintained consistently high agreement with the reference clustering as *λ* increased, indicating strong stability to between-sample variation for this dataset. In contrast, some methods showed pronounced degradation under stronger variation (e.g., GMM), while others exhibited relatively low stability even at small *λ* (e.g., Ward hierarchical clustering). Given the strong stability of Louvain clustering for this dataset, we next used scStable to select stable hyperparameters within the Louvain workflow.

We focused on the standard Seurat Louvain workflow, controlled by three key hyperparameters: the number of input principal components (*n*_PC_), the number of nearest neighbors in the shared nearest neighbor graph (*k*_NN_), and the Louvain resolution (*r*). These hyperparameters are commonly chosen by visual inspection of clustering results or set to software defaults. We constructed a grid of hyperparameter combinations within typical ranges and used scStable synthetic scRNA-seq samples to evaluate the stability of each combination. For each combination, we measured agreement between the clustering from each synthetic scRNA-seq sample and that from the single-sample scRNA-seq reference at *λ* = 1 using the four agreement metrics described above (Methods; complete results in Supplementary Fig. S4). The combination *n*_PC_ = 10, *k*_NN_ = 20, and *r* = 0.40 achieved the highest agreement across synthetic scRNA-seq samples and was therefore selected for this dataset.

Fig. 5C compares the clusters obtained under the default and scStable-selected hyperparameters, with reference cell-type labels used only for evaluation. Under the default hyperparameters, the B cell population was split into two clusters, and regulatory T cells were mixed with neighboring naive cytotoxic cells. With the scStable-selected hyperparameters, the clusters agreed more closely with the reference cell types, with ARI increasing from 0.763 to 0.899. Thus, for this dataset, selecting hyperparameters based on stability across synthetic scRNA-seq samples improved agreement with the reference cell-type labels relative to the default hyperparameters.

## Discussion

scStable addresses a common limitation of scRNA-seq studies in which only one or a few biological samples are available: the inability to assess how sensitive computational discoveries are to between-sample variation. By transferring bulk-derived between-sample variation to a generative model fitted to a single-sample scRNA-seq reference while preserving cell-level correspondence across synthetic scRNA-seq samples, scStable enables stability-aware analysis without requiring additional scRNA-seq samples. Our results illustrate two complementary uses of this framework: assessing the stability of downstream discoveries, such as DE genes, and stability-driven, data-specific selection of methods and hyperparameters, such as for cell clustering. The analysis of paired multi-sample scRNA-seq and bulk RNA-seq data from gastric cancer further showed that the between-sample variation generated by scStable more closely recapitulated that observed in held-out real multi-sample scRNA-seq data than did ZINB-WaVE, Gaussian noise injection, or cell-wise subsampling.

The stability assessed by scStable should be interpreted as robustness to the modeled bulk-derived between-sample variation, rather than as a guarantee that a result will generalize to future biological samples. Stability is also not a strict requirement for biological relevance: signals that are sensitive to between-sample variation, including potentially rare or context-specific signals, may still be scientifically important. Instead, scStable makes this sensitivity explicit, allowing stability to be considered alongside statistical evidence and biological knowledge when prioritizing discoveries or selecting analysis procedures. Similarly, instability under scStable indicates sensitivity to the modeled variation rather than evidence that a result is incorrect.

A practical consideration is the choice of multi-sample bulk RNA-seq reference. In bulk-reference mode, bulk RNA-seq from the same study or a closely matched cohort can provide condition- and tissue-specific between-sample variation. When such data are unavailable, bulk-reference-free mode can use public tissue-matched bulk RNA-seq resources such as GTEx. The quality of the resulting stability assessment therefore depends on how well the bulk reference represents the biological population relevant to the single-sample scRNA-seq reference. The scale factor *λ* provides an additional way to vary the magnitude of injected between-sample variation: *λ* = 1 retains the variation estimated from the bulk reference, whereas larger values can be used for stress testing. Our gastric cancer analysis suggests that the unamplified bulk-derived variation provides a reasonable default for that dataset, although the appropriate *λ* may depend on the application.

An important limitation is that scStable preserves the cells and therefore the cell-type composition of the single-sample scRNA-seq reference. In contrast, between-sample variation in bulk RNA-seq reflects both within-cell-type expression changes and changes in cell-type composition. Because bulk RNA-seq does not directly separate these components, transferring bulk-derived variation to a fixed set of cells cannot be expected to reproduce all aspects of real multi-sample scRNA-seq variation simultaneously. This limitation was reflected in the gastric cancer analysis, where pseudo-bulk- and single-cell-level agreement with real scRNA-seq samples favored somewhat different values of *λ*. Incorporating cell-type-aware or deconvolution-based modeling could help distinguish expression and compositional variation, although doing so reliably from bulk RNA-seq introduces additional uncertainty.

Several extensions could further broaden the scope of scStable. Curated libraries of pre-estimated, tissue- and condition-specific between-sample variation profiles could make bulk-reference-free analysis more readily applicable when well-matched bulk RNA-seq data are unavailable. The same general principle of combining a single-sample generative model with empirically learned between-sample variation could also be extended to spatial transcriptomics and other single-cell omics modalities, provided that modality-specific structure can be preserved while sample-level variation is introduced.

## Conclusions

scStable provides a framework for assessing the stability of scRNA-seq analyses when biological replicates are limited or unavailable by generating synthetic scRNA-seq samples that preserve cell-level heterogeneity while incorporating bulk-derived between-sample variation. By enabling both stability-aware assessment of downstream discoveries and stability-driven, data-specific selection of analysis methods and hyperparameters, scStable complements conventional single-sample scRNA-seq analysis with an explicit assessment of sensitivity to between-sample variation.

## Methods

### Datasets and preprocessing

The GTEx whole-blood bulk RNA-seq dataset [27] includes 13,807 genes across 407 samples, obtained from the GTEx Portal open-access downloads.

The Zheng4 dataset [29] is a 10x Genomics UMI-based scRNA-seq mixture of four purified human peripheral blood mononuclear cell (PBMC) populations with known labels, containing 3,994 cells and 15,568 genes. We used the `Zhengmix4eq` version curated by Duò et al. [34], in which the four populations are represented in approximately equal proportions, obtained through the Bioconductor experiment data package `DuoClustering2018` [35].

The aging mouse lung dataset [32] provides Drop-seq scRNA-seq of whole lung tissue from young (3 months) and old (24 months) mice, together with bulk RNA-seq of flow-sorted lung cell populations. For our analysis, we subset macrophage cells from the scRNA-seq data, retaining 17,681 genes and 1,183 cells from young mice and 521 cells from old mice, and used flow-sorted macrophage bulk RNA-seq as the corresponding multi-sample bulk reference (young *n* = 5, old *n* = 7; *∼*30,000 genes). The lung aging analysis was intended as a proof-of-concept demonstration of condition-level stability assessment within a single cell type. Because individual mice contributed too few macrophage cells for stable model fitting after restricting the analysis to this cell type, we pooled macrophage cells across mice within each age group to construct age-specific pooled scRNA-seq references, containing 1,183 cells for young mice and 521 cells for old mice. The mice were from the same inbred strain, reducing biological heterogeneity across animals. Accordingly, this analysis demonstrates the condition-level stability workflow rather than a literal single-biological-sample application. Data and metadata were accessed through GEO (accession `GSE124872`).

The paired gastric tumor microenvironment dataset [30] contains matched malignant and adjacent non-malignant gastric tissue from 24 treatment-naive gastric cancer patients, profiled using both 10x Genomics scRNA-seq and bulk RNA-seq. The published single-cell analysis retained 96,623 cells after quality control and downstream filtering. Processed data were obtained from the dataset released by the original authors; the corresponding GEO accession is `GSE206785`. The processed scRNA-seq and matched bulk RNA-seq data used in our analysis were obtained from the original authors’ public GitHub repository (https://github.com/chriscainx/gastric-cancer). For our comparison analysis, we restricted the dataset to non-malignant gastric tissue and excluded scRNA-seq samples containing fewer than 100 cells, leaving 21 patients with matched scRNA-seq and bulk RNA-seq profiles. One scRNA-seq sample was designated as the single-sample scRNA-seq reference, and the remaining 20 scRNA-seq samples constituted the real multi-sample scRNA-seq comparison set; the corresponding 21 bulk RNA-seq profiles were used as the multi-sample bulk RNA-seq reference.

Across all datasets, genes were filtered jointly on the multi-sample bulk and single-sample scRNA-seq references before generation. A gene was retained if fewer than 20% of bulk samples had expression equal to the gene-specific minimum and if the proportion of zero counts across single cells was below 95%. Only genes passing both criteria and detected in at least 2 cells were kept for downstream analysis; no additional cell-level filtering was applied.

### scStable generation workflow

scStable is designed to generate synthetic scRNA-seq samples from a single-sample scRNA-seq reference by preserving cell-level heterogeneity from the reference while injecting bulk-derived between-sample variation learned from a multi-sample bulk RNA-seq reference, either from the same study as the single-sample scRNA-seq reference (bulk-reference mode) or from a public tissue-matched resource (bulk-reference-free mode). The generation workflow consists of four steps: fitting a cell-label-free scRNA-seq generative model, estimating bulk-derived between-sample variation, mapping this variation to the single-cell generative model, and generating synthetic scRNA-seq samples for stability assessment and method selection.

#### Step 1: Model single-sample scRNA-seq reference

The goal of this step is to fit *a scRNA-seq generative model* that captures cell-level heterogeneity from the single-sample scRNA-seq reference. scStable builds on scDesign3 [21], a state-of-the-art simulator that incorporates cellular covariates to preserve cell-level heterogeneity. To support the generation of synthetic scRNA-seq samples with between-sample variation learned from the multi-sample bulk RNA-seq reference in later steps, scStable extends scDesign3 in two ways: (i)by adopting a cell-label-free modeling strategy that preserves cell-level heterogeneity without requiring cell type/state or trajectory annotations, enabling data-specific stability evaluation for cell-level analyses (e.g., clustering and trajectory inference) where such labels are unavailable, and (ii)by enabling sample-specific adjustments of fitted model parameters informed by bulk-derived between-sample variation (Step 3).

Let 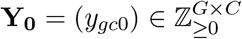 denote the gene-by-cell count matrix of the single-sample scRNA-seq reference (indexed by “0”), where rows correspond to *G* genes, and columns correspond to *C* cells. Following scDesign3, we first model each gene *g* = 1, …, *G* using a marginal (per-gene) count distribution:

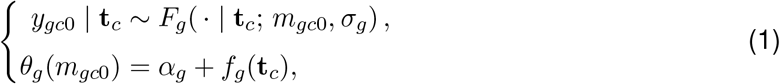

where **t**_*c*_ denotes a vector of cell-specific covariates for cell *c* = 1, …, *C*, used to capture cell-level heterogeneity. Importantly, **t**_*c*_ is not indexed by sample, because scStable generates synthetic scRNA-seq samples by using the same set of cells (and their covariates) as in the single-sample scRNA-seq reference. Here, *F*_*g*_ denotes the count distribution for gene *g*, with *m*_*gc*0_ as the gene- and cell-specific mean parameter for single-sample scRNA-seq reference and *σ*_*g*_ as a gene-specific scale parameter (e.g., dispersion for the negative binomial distribution). The function *θ*_*g*_(*·*) is a gene-specific link function relating *m*_*gc*0_ to the covariates **t**_*c*_ through *f*_*g*_(**t**_*c*_), with *α*_*g*_ denoting a gene-specific intercept.

By default, we use the negative binomial distribution for *F*_*g*_ and the log link function *θ*_*g*_(*·*) = log(*·*) [36]. The covariates **t**_*c*_ are chosen as the top *k* principal components (PCs) of the single-sample scRNA-seq reference, serving as a surrogate for cell type or cell state, with *k* user-specified (default *k* = 10 based on sensitivity analysis). We model *f*_*g*_(*·*) using cubic spline functions of the PCs.

Beyond marginal per-gene modeling, we next account for gene–gene dependence using the same Gaussian copula-based joint modeling strategy as in scDesign3. Specifically, each gene’s counts are transformed to Gaussian variables based on the fitted marginal distributions, and a multivariate Gaussian model is used to capture second-order gene–gene dependencies via a gene-gene correlation matrix. Model fitting yields estimates of the gene-by-cell mean matrix 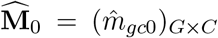 for the single-sample scRNA-seq reference, along with additional estimated parameters, including gene-specific dispersions and the Gaussian copula correlation matrix. In Step 3, scStable injects between-sample variation learned in Step 2 by adjusting 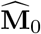 in a sample-specific manner, while keeping the remaining estimated parameters fixed during sample generation.

#### Step 2: Model multi-sample bulk RNA-seq reference

The goal of this step is to learn a distribution that captures *between-sample variation* from a multi-sample bulk RNA-seq reference, which will later be mapped to the scRNA-seq generative model. In bulk-reference mode, we assume that the supplied bulk RNA-seq reference represents the same or a closely matched tissue, biological condition, and study population as the single-sample scRNA-seq reference. In bulk-reference-free mode, the public reference is tissue-matched but need not be study- or condition-matched. Under these assumptions, variation across bulk samples provides an empirical template for between-sample variation relevant to the scRNA-seq reference. To ensure compatibility between modalities (single-cell vs. bulk), scStable learns gene-specific variation using the set of genes shared between the single-sample scRNA-seq reference and the multi-sample bulk RNA-seq reference. The procedure below is the same in both modes; the modes differ only in whether the multi-sample bulk RNA-seq reference is study-matched or public, as described at the end of this section.

Let 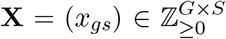 denote the bulk RNA-seq count matrix, where rows correspond to the *G* genes retained after filtering (Datasets and preprocessing) and columns correspond to *S* bulk samples. We normalize each sample to counts per million (CPM),

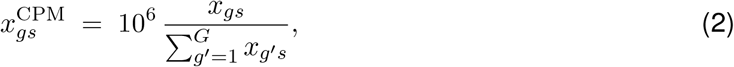

and denote the resulting matrix by 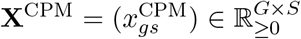. Motivated by the widespread use of log-CPM in bulk RNA-seq analysis and Gaussian modeling of transformed expression values [37], we apply a gene-specific log transformation to the CPM values,

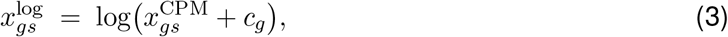

where *c*_*g*_ *>* 0 is a gene-specific pseudo-count. The value of *c*_*g*_ is chosen to make the transformed expression values 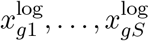 approximately Gaussian, as assessed by a normality test (see Supplementary Section S1). Let 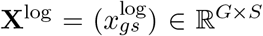 denote the resulting log-CPM expression matrix, whose *s*-th column 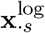 represents the sample *s*, a *G*-dimensional gene expression vector.

After the transformation, we assume

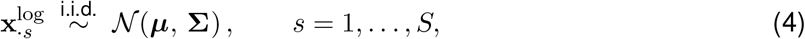

where ***µ*** ∈ ℝ^*G*^ and **S** ∈ ℝ^*G×G*^ represent the mean vector and covariance matrix of *G* genes, capturing between-sample variation. Because the number of genes *G* can be large relative to the number of bulk samples *S*, we estimate **Σ** using a regularized covariance estimator when *S ≥* 30 [38], and use a diagonal approximation when *S <* 30 (to ensure computational stability and avoid overfitting). The fitted parameters 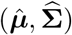 characterize bulk-derived between-sample variation and serve as input to Step 3.

The two modes differ only in the source of the multi-sample bulk RNA-seq reference. In bulk-reference mode, it comes from the same study as the single-sample scRNA-seq reference, profiled either from the same individuals or from an independent cohort of the same population. In bulk-reference-free mode, no study-matched bulk RNA-seq is available, and the reference is public tissue-matched bulk RNA-seq such as GTEx [27]. The user either supplies the public count matrix, to which the gene filtering (Datasets and preprocessing), the pseudo-count selection and the estimation of 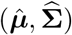 above are applied unchanged, or specifies a tissue name, in which case scStable uses 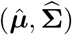 estimated from the corresponding GTEx data by the same procedure.

In addition, scStable introduces a user-adjustable scale factor in Step 3 to modulate the magnitude of injected between-sample variation. This scaling mechanism allows users to calibrate the variation level when default bulk-derived between-sample variation may not fully match their study population, and enables stress-testing of downstream analyses across a range of between-sample variation magnitudes.

#### Step 3: Map bulk-derived between-sample variation

The goal of this step is to map bulk-derived between-sample variation, learned in Step 2, onto the scRNA-seq generative model fitted in Step 1. This mapping consists of three operations: (i) learning gene-specific offsets on the log-CPM scale to align the reference scRNA-seq pseudo-bulk with the bulk RNA-seq samples; (ii) generating *R* synthetic bulk samples from (4) and converting them into pseudo-bulk samples using these offsets; and (iii) using the *R* synthetic pseudo-bulk samples together with the single-sample scRNA-seq reference pseudo-bulk to compute *R* sets of per-gene multiplicative adjustments, which are applied to the mean matrix of the scRNA-seq generative model to obtain *R* adjusted models.

Because scRNA-seq data and bulk RNA-seq data are on different scales, we first align the pseudo-bulk of the single-sample scRNA-seq reference to the multi-sample bulk RNA-seq reference. We define the pseudo-bulk count for gene *g* from the single-sample scRNA-seq reference as

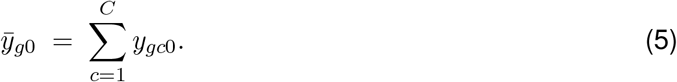

We normalize the pseudo-bulk to CPM over the same *G* genes as in (2), 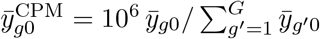 and apply the same gene-specific log transformation used for the multi-sample bulk RNA-seq reference as in (3),

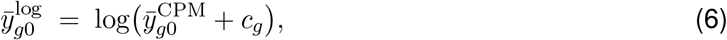

where *c*_*g*_ is the gene-specific pseudo-count determined in Step 2.

To align the pseudo-bulk of the single-sample scRNA-seq reference with the multi-sample bulk RNA-seq reference on the log-CPM scale (to ensure that sample library size does not confound the alignment), we introduce a gene-specific offset

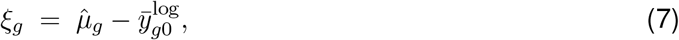

where 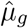 is the *g*-th component of the estimated bulk mean vector 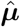 from Step 2. This offset accounts for gene-specific differences in mean log-CPM expression between the scRNA-seq pseudo-bulk and the multi-sample bulk RNA-seq reference. We assume that modality differences can be approximated by a per-gene location shift on the log-CPM scale, allowing deviations around the bulk mean to be transferred to the scRNA-seq reference pseudo-bulk. Under this assumption, 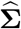 characterizes the covariance of between-sample deviations across genes, while *ξ*_*g*_ aligns the mean expression level of each gene across the two modalities.

Using this alignment, we generate *R* synthetic scRNA-seq samples by constructing *R* sample-specific variants of the scRNA-seq generative model. For each synthetic scRNA-seq sample *r* = 1, …, *R*, we first draw a bulk-level expression vector on the log-CPM scale from the fitted bulk Gaussian model in Step 2, restricted to the region on which the inverse transformation below is non-negative,

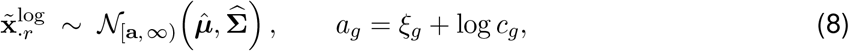

where *N*_[**a**, *∞*)_ denotes the Gaussian of (4) truncated to 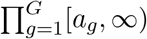, with *a*_*g*_ denoting the gene-specific lower bound corresponding to 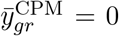 under (9). The truncation thus guarantees that the pseudo-bulk CPM values obtained below, and hence the multiplicative adjustments *ω*_*gr*_ in (10), are non-negative by construction. When 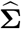 is diagonal (*S <* 30), the truncated Gaussian factorizes into *G* independent univariate truncated Gaussians.

We then map 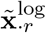 back to the pseudo-bulk CPM scale by removing the offset and inverting the log transformation,

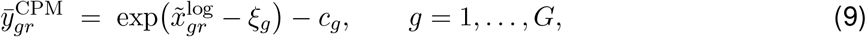

where 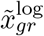 is the *g*-th component of the vector 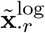. By (8), 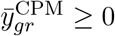 for all *g* and *r*.

Next, we inject between-sample variation into the scRNA-seq model by applying a per-gene multiplicative adjustment to the estimated mean matrix 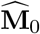 from Step 1. Specifically, for each synthetic scRNA-seq sample *r*, we define

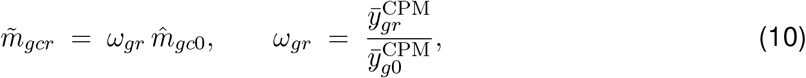

for *g* = 1, …, *G* and *c* = 1, …, *C*. This per-gene scaling applies a shared sample-specific gene expression shift *ω*_*gr*_ across all cells for each gene, reflecting bulk-derived between-sample variation while preserving the cell-level heterogeneity encoded in 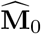. The resulting mean matrix for synthetic scRNA-seq sample *r* is denoted by 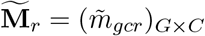.

To further control the magnitude of injected between-sample variation, scStable introduces a user-adjustable scale factor *λ*. This parameter interpolates between no injected variation (*λ* = 0), the bulk-derived between-sample variation (*λ* = 1), and amplified variation (*λ >* 1) for stress-testing downstream analyses. Specifically, Eq. (10) is modified to

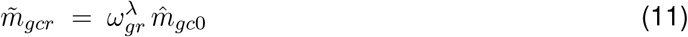

At *λ* = 0, we define *ω*^0^ = 1, so that no between-sample variation is injected.

#### Step 4: Generate synthetic scRNA-seq samples

To generate the *r*-th synthetic scRNA-seq sample, we use the adjusted gene-by-cell mean matrix 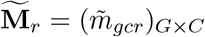 obtained in Step 3, together with the remaining fitted parameters of the scRNA-seq generative model from Step 1. Specifically, synthetic scRNA-seq samples are generated by sampling from the same generative model as in Step 1, with the estimated mean matrix 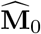 replaced by 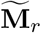. The sampling procedure follows scDesign3: we first generate latent Gaussian variables using the fitted copula model to capture gene–gene dependence, and then transform them into per-gene counts according to the marginal distributions in (1), using 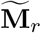 as the mean parameters.

This procedure preserves the cell-level heterogeneity encoded in the single-sample scRNA-seq reference while introducing sample-specific variation governed by bulk-derived between-sample variation. The bulk model in Step 2 captures cross-gene covariance of sample-level expression through 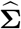 (or only per-gene variances when a diagonal approximation is used), but it does not identify gene–gene dependence across cells within an scRNA-seq sample. Accordingly, the cell-level gene–gene dependence structure estimated from the single-sample scRNA-seq reference in Step 1 is held fixed across synthetic scRNA-seq samples.

### Downstream analysis methods

We apply standard scRNA-seq analysis pipelines to the single-sample scRNA-seq reference and to each synthetic scRNA-seq sample produced by scStable.

#### Differential expression

For between-cell-type and between-condition DE analyses on individual samples, we use the six DE methods in Seurat v5 [31]: the Wilcoxon rank-sum test, negative binomial and Bimod likelihood ratio tests, Student’s *t*-test, logistic regression, and Poisson likelihood ratio test. Raw *p*-values are adjusted for multiple testing across all tested genes using the Benjamini–Hochberg (BH) procedure. The resulting BH-adjusted *p*-value, denoted *p*_adj_, serves as the scalar summary *T* used by the stability metrics defined below. For between-condition DE (lung aging dataset), the *r*-th young and *r*-th old synthetic scRNA-seq samples are combined into the *r*-th synthetic two-condition dataset, *r* = 1, …, *R*, and the Wilcoxon rank-sum test is applied to the young-versus-old comparison in each, as in the reference.

#### Clustering

We evaluate six clustering methods: Louvain, Louvain with multi-level refinement, and Smart Local Moving (SLM) from Seurat v5 [31]; *K*-means and Ward hierarchical clustering in base R; and Gaussian mixture modeling from `mclust` [33]. Unless otherwise noted, Seurat clustering uses the top 10 PCs and a *k*-nearest-neighbor graph with *k* = 20 at resolution 0.8. *K*-means (10 random starts), Ward hierarchical clustering, and the Gaussian mixture model require the number of clusters to be specified, and we set *K* = 4, the number of cell populations in Zheng4. For these three methods, we set *K* = 4 to match the four known cell populations in Zheng4; in applications without known labels, *K* would be user-specified or selected by the analyst. For the joint hyperparameter analysis, we vary *n*_PC_ ∈ *{*10, 20, 50*}, k*_NN_ ∈ *{*10, 20, 50*}*, and the Louvain resolution *r* ∈ *{*0.4, 0.8, 1.2*}*.

#### Pathway enrichment

KEGG pathway enrichment is performed separately for stable DE genes and other DE genes using `clusterProfiler` [39], with all genes tested for DE used as the background gene universe. Species-matched KEGG databases are used for each analysis (human for Zheng4 and mouse for the lung aging dataset), and *p*-values are adjusted using the Benjamini–Hochberg procedure. For each gene set, we report the top pathways ranked by adjusted *p*-value together with the corresponding gene ratio.

#### Agreement metrics and embeddings

We measure agreement between two clusterings of the same *C* cells using four standard metrics. Let *a* denote the number of cell pairs that are placed in the same cluster by both clusterings, and let *b* and *c* denote the numbers of pairs that are placed together by only the first or only the second clustering, respectively. From these pair counts we compute the Jaccard index, *a/*(*a* + *b* + *c*), and the Fowlkes–Mallows index (FMI), 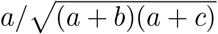. We also report the Adjusted Rand Index (ARI), which corrects the pair-counting Rand index for agreement expected by chance [40], and the Normalized Mutual Information (NMI) [41]. All four metrics are invariant to permutations of cluster labels and equal 1 when the two clusterings are identical, so they can be applied directly to clusterings obtained from different samples without matching cluster labels.

We quantify how well cells with different labels are mixed in an embedding using the local inverse Simpson’s index (LISI) [42]. For a categorical label with *J* levels, the LISI of cell *i* is defined as

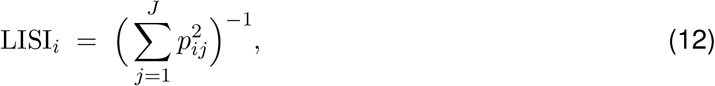

where *p*_*ij*_ is the proportion of cell *i*’s neighbours that carry level *j*, with neighbours weighted by a Gaussian kernel whose bandwidth corresponds to a perplexity of 30. LISI_*i*_ can be interpreted as the effective number of levels represented among the neighbours of cell *i*; it equals 1 when all neighbours share a single level and *J* when all levels are equally represented. We summarize LISI across cells by its median, denoted mLISI. In the validation analysis, the label distinguishes reference cells from synthetic cells (*J* = 2), and mLISI is computed on the two-dimensional UMAP coordinates described below, so that values near 2 indicate that synthetic cells are locally indistinguishable from reference cells, whereas values near 1 indicate that the two occupy separate neighbourhoods. In the comparison analysis, the label is sample identity (*J* = 20), as described in Single-cell level: variation metrics.

To visualize synthetic scRNA-seq samples in the coordinates of the single-sample scRNA-seq reference (Figs. 2A and 4E), we fit principal component analysis (PCA) on the centered and scaled log(1 + *x*) counts of the reference, retaining the top 20 PCs, and fit UMAP on these PCs using the default parameters of the `umap` R package. Synthetic cells are then projected onto the reference PC basis and mapped through the fitted UMAP model. Because the embedding is fitted on the reference alone, its coordinates are fixed across panels, so that differences between panels reflect differences in the synthetic data rather than changes in the embedding.

### Stability assessment and method selection

The output of the scStable generation workflow is a collection of synthetic scRNA-seq samples 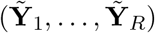, with each 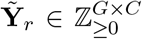 generated with bulk-derived between-sample variation (Steps 1–4). The metrics below define how these samples are used for stability-aware assessment of downstream discoveries and stability-driven method selection.

#### Quantile-interval stability for downstream discoveries

When only a single-sample scRNA-seq reference **Y**_0_ is available, a downstream pipeline yields a single realization of the result (e.g., a DE statistic, a trajectory, or a cell–cell communication network). Internal uncertainty measures such as *p*-values do not by themselves quantify sensitivity to between-sample variation that could arise across biologically similar samples. scStable instead enables a sample-based assessment by applying the same procedure independently to each synthetic scRNA-seq sample 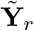.

Let *A* denote an analysis procedure and *T* (*·*) a scalar summary of its output (e.g., a per-gene adjusted *p*-value, log-fold change, or edge weight). For each synthetic scRNA-seq sample *r*, we compute

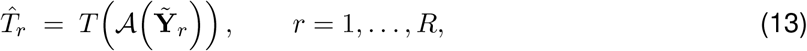

yielding an empirical distribution 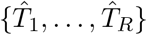 that captures variability induced by between-sample variation. We summarize it by a (1 *− α*) quantile interval across synthetic scRNA-seq samples,

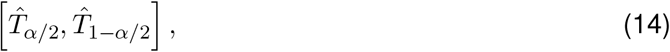

where 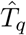 is the *q*-th empirical quantile of 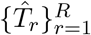, and define stability by the interval length 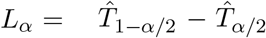, with shorter intervals indicating greater stability. Throughout, we set *α* = 0.05 and call a gene a *stable DE gene* if it is significant in the single-sample scRNA-seq reference (*p*_adj_ *<* 0.05) and satisfies *L*_*α*_ *<* 0.05. Genes that are significant in the reference but do not meet the stability criterion are referred to as *other DE genes*.

#### Entropy-based per-cell clustering stability

To assess clustering stability in a label-agnostic manner (cluster labels are arbitrary across independent samples), we use a consensus matrix approach [41, 43]. For each synthetic scRNA-seq sample *r* = 1, …, *R*, we cluster the cells to obtain labels 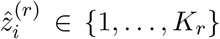 and form a binary cell–cell co-clustering matrix **M**^(*r*)^ ∈ *{*0, 1*}*^*C×C*^,

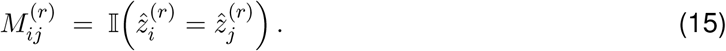

Averaging over the *R* samples gives the consensus matrix **C** ∈ [0, 1]^*C×C*^,

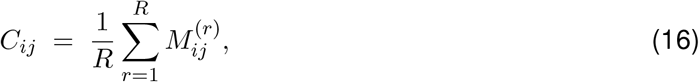

where *C*_*ij*_ near 1 indicates consistent co-clustering, near 0 consistent separation, and near 0.5 an unstable relationship. We then score per-cell stability by treating each *C*_*ij*_ as a Bernoulli success probability and computing its binary entropy,

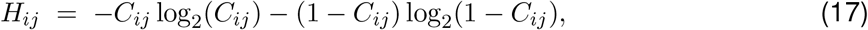

with *H*_*ij*_ = 0 when *C*_*ij*_ ∈ *{*0, 1*}* (decisive) and *H*_*ij*_ = 1 when *C*_*ij*_ = 0.5 (maximally uncertain). Averaging over all other cells gives

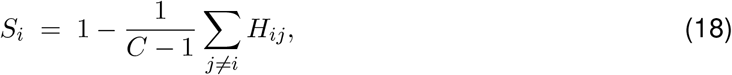

with larger *S*_*i*_ indicating more decisive, stable cluster assignments under between-sample variation.

#### Stability-driven method selection

Let *A*_1_, …, *A*_*L*_ denote *L* candidate analysis specifications for a fixed task, where a specification may be a distinct method or a particular hyperparameter setting. Given synthetic scRNA-seq samples 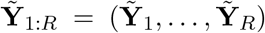, we apply each specification *A*_*ℓ*_ to every synthetic scRNA-seq sample and to the single-sample scRNA-seq reference **Y**_0_, and define its stability as the average agreement between the two outputs,

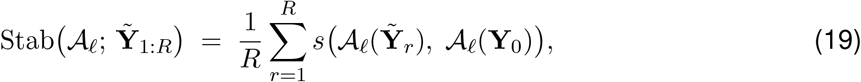

where *s*(*·, ·*) is a task-appropriate similarity between two outputs, with larger values indicating greater stability. Because every synthetic scRNA-seq sample contains the same cells as **Y**_0_ (Step 4), outputs defined on cells can be compared cell by cell. For clustering, *A*_*ℓ*_(*·*) is a vector of cluster labels over the *C* cells and *s* is one of the four agreement metrics (ARI, NMI, Jaccard index, FMI; Agreement metrics and embeddings), each computed separately; these metrics are invariant to permutation of cluster labels, so no label matching across samples is required. To stress-test specifications, we generate synthetic scRNA-seq samples 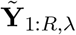 at each value of the scale factor *λ* (Step 3) in a grid Λ and evaluate each specification along this variation path,

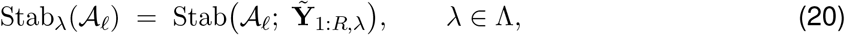

with Λ = *{*0, 0.05, 0.1, 0.2, 0.3, 0.5, 0.7, 1.0, 1.5, 2.0*}* and *R* = 100 synthetic scRNA-seq samples per *λ* in this work (Supplementary Table S1).

The two uses of Eq. (19) differ only in the candidate set and the *λ* values considered. For method selection, the candidates are distinct methods, and we compare the curves *λ 1→* Stab_*λ*_(*A*_*ℓ*_) over Λ, preferring methods whose stability remains high as *λ* increases over methods whose stability is low at small *λ* or degrades as *λ* increases. For hyperparameter selection within a fixed method, each candidate hyperparameter vector (for Louvain clustering, the number of principal components *n*_PC_, the number of nearest neighbors *k*_NN_, and the resolution) is treated as a separate specification, and we select the setting that maximizes Stab_1_(*A*_*ℓ*_), i.e., stability at the bulk-derived level of between-sample variation, across the four agreement metrics. Cell-type labels are not used at any point in either selection.

### Comparing against alternative approaches

To compare scStable with alternative strategies for generating synthetic scRNA-seq samples from a single-sample scRNA-seq reference, we compare four generators on the paired gastric tumor microenvironment dataset: (i) scStable; (ii) ZINB-WaVE [44], a representative model-based single-sample simulator; (iii) Gaussian noise injection; and (iv) cell-wise subsampling. Real multi-sample scRNA-seq from the same study provides the empirical reference for the comparison, while the paired bulk RNA-seq serves as an additional empirical anchor. Sample 180118N is held out as the single-sample scRNA-seq reference.

#### scStable

Synthetic scRNA-seq samples are generated as described in scStable generation workflow Section, with the scale factor *λ* controlling the magnitude of bulk-derived between-sample variation injected into each synthetic scRNA-seq sample. We generate synthetic scRNA-seq samples at each of *λ* ∈ *{*1, 1.6, 1.7, 1.8, 2*}*.

#### ZINB-WaVE

We selected ZINB-WaVE as a representative model-based single-sample simulator because, in our implementation, it permits one-to-one cell correspondence with the reference, enabling direct per-cell comparisons while retaining a common gene set. ZINB-WaVE has also shown competitive performance in previous evaluations of scRNA-seq simulators [22, 28]. We fit ZINB-WaVE [44] to the single-sample scRNA-seq reference using two latent factors and generate synthetic scRNA-seq samples from the fitted model while retaining the fitted cell-specific parameters, so that each simulated cell corresponds one-to-one to a cell in the reference. Across synthetic scRNA-seq samples, the cells and fitted parameters are held fixed, and samples differ only by sampling stochasticity, with no injected between-sample variation.

#### Gaussian noise injection

This baseline adds gene-specific noise directly to the single-sample scRNA-seq reference and uses no bulk RNA-seq data. All operations are on the `log` scale, which keeps the added noise multiplicative on the count scale. For each gene *g* we take the standard deviation of log(1 + *y*_*gc*0_) across the *C* cells of the reference and cap it at the 95th percentile of that distribution over genes,

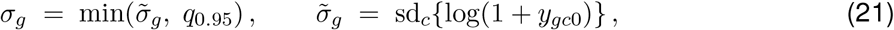

where *q*_0.95_ is the 95th percentile of 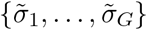; the cap prevents a few genes expressed highly in a minority of cells from dominating. For each synthetic scRNA-seq sample *r* we draw one offset per gene, independently across genes and samples, and add it to every cell of the reference,

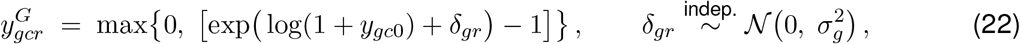

where [*·*] denotes rounding to the nearest integer.

#### Cell-wise subsampling

For each synthetic scRNA-seq sample we draw *C*^*′*^ = 0.8 *C* cells without replacement from the single-sample scRNA-seq reference, and rescale the resulting pseudo-bulk profile by 1*/*0.8 to restore the reference sequencing depth. This captures within-sample sampling variability but introduces no biological between-sample variation.

#### Real multi-sample scRNA-seq

From the paired gastric tumor microenvironment dataset [30], we designated Sample 180118N as the single-sample scRNA-seq reference and used the remaining normal-tissue samples containing at least 100 cells as the real multi-sample scRNA-seq comparison set, yielding *n* = 20 samples. These samples are used only for comparison and are never used during synthetic sample generation.

#### Pseudo-bulk construction and normalization

All pseudo-bulk comparisons are made on the log-CPM scale introduced in Step 2, so that profiles derived from scRNA-seq and from bulk RNA-seq are directly comparable. Extending Eq. (5) from the reference to an arbitrary sample *s*, the pseudo-bulk count of gene *g* is

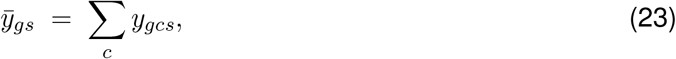

summed over the cells of that sample, where *s* indexes a sample from the real multi-sample scRNA-seq or a synthetic scRNA-seq sample.

Because Eq. (23) is a plain sum, a sample’s total scales with its cell number. Real multi-sample scRNA-seq samples differ several-fold in cell number, whereas every synthetic scRNA-seq sample has the *C* cells of the single-sample scRNA-seq reference, so on the raw-sum scale the apparent between-sample variation is strongly driven by differences in captured cell number and is therefore not directly comparable with synthetic samples that have a fixed cell count. The real samples contain between 202 and 10,658 cells, and on the raw-sum scale their leading principal component accounts for 87% of the total variance and correlates with log cell number at *r* = *−*0.94. We therefore process every pseudo-bulk profile exactly as the reference pseudo-bulk in Step 3: we normalize it to CPM as in Eq. (2),

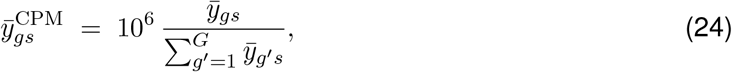

apply the log transformation of Eq. (6), and add the gene-specific offset *ξ*_*g*_ of Eq. (7),

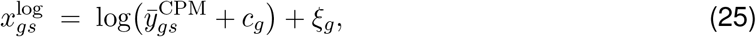

with *c*_*g*_ the gene-specific pseudo-count from Step 2. Paired bulk RNA-seq is already on this scale by Eqs. (2)–(3). All results in the main text use these library-size-normalized pseudo-bulk profiles. Without this normalization the separation between scStable and the alternatives is unchanged.

For descriptive comparison, we used a common panel of 1,369 genes after excluding any gene whose between-sample variance fell in the top 10% in at least one compared group (real multi-sample scRNA-seq, paired bulk RNA-seq, or any synthetic approach). The same resulting gene panel was then used for all groups. This symmetric trimming rule prevents a small number of extreme variances from dominating the descriptive comparisons and was not used for hypothesis testing or statistical significance assessment.

#### Per-gene between-sample mean and variance

For a group of *S* samples (real multi-sample scRNA-seq, paired bulk RNA-seq, or one generator setting), the per-gene mean and the per-gene between-sample variance are

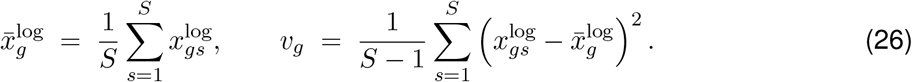

#### Sensitivity to the scale factor *λ*

For each *λ* in the grid we compute 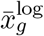 and *v*_*g*_ over the synthetic scRNA-seq samples and compare them with the corresponding quantities from real multi-sample scRNA-seq and paired bulk RNA-seq. Three statistics summarize the comparison. First, the Spearman rank correlation of 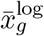 and of *v*_*g*_ against their real counterparts, which tests whether the *ordering* of genes is preserved. Second, Lin’s concordance correlation coefficient [45] between a synthetic statistic *u* and its real counterpart *w*,

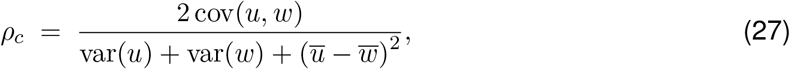

which measures agreement with the identity line *y* = *x* rather than mere association; unlike the Spearman and Pearson correlations it is not invariant to rescaling one argument, so a setting scores highly only by producing variances of the correct magnitude as well as the correct ordering. It is evaluated on the scale on which each quantity is displayed: linearly for 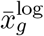, and on log_10_ for *v*_*g*_ (genes with *v*_*g*_ = 0 in either group are excluded from this statistic only). Third, the total between-sample variance ratio

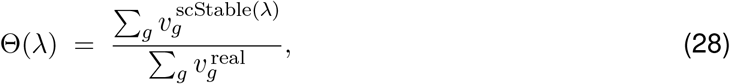

where Θ = 1 indicates that the synthetic scRNA-seq samples carry exactly as much total between-sample variation as the real multi-sample scRNA-seq. We use Θ(*λ*) to describe how the total between-sample variation responds to *λ*, not to select a single value; *ρ*_*c*_ and the Spearman correlations likewise describe what *λ* does and does not change. Uncertainty in Θ comes from resampling genes with replacement 2,000 times and reporting the 2.5th and 97.5th percentiles. Θ is also reported against paired bulk RNA-seq; the two references are approached from opposite directions. The full curve is given in Supplementary Fig. S1.

#### Between-sample variation metrics

Three complementary metrics compare each approach with real multi-sample scRNA-seq, showing *which* genes vary between samples and *how much* they vary. All are computed on the library-size-normalized common gene panel, with real multi-sample scRNA-seq as the reference group and 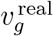 its per-gene between-sample variance.

The *median variance ratio* is the per-gene ratio of between-sample variance to real multi-sample scRNA-seq, summarized across genes by its median,

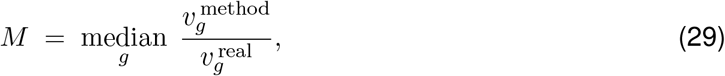

and reported on a log_10_ axis (Fig. 3C). *M* = 1 indicates that a typical gene carries exactly the real amount of between-sample variation; *M* = 0.5 indicates half as much.

The *total variance ratio* is the ratio Θ of Eq. (28), evaluated for each approach rather than each *λ* (Fig. 3D). Unlike the median variance ratio, which describes a typical gene, it describes the aggregate amount of between-sample variation and is therefore dominated by the genes that vary most.

*Concordance* is Lin’s concordance correlation coefficient *ρ*_*c*_ of Eq. (27) between 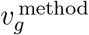 and 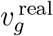 across genes, evaluated on the log_10_ scale and excluding genes with *v*_*g*_ = 0 in either group (Fig. 3E). Because *ρ*_*c*_ is not invariant to rescaling, it rewards agreement in both the ordering and the magnitude of per-gene variance.

#### Single-cell level: preprocessing

We repeat the comparison at the single-cell level. Real multi-sample scRNA-seq samples contain between 202 and 10,658 cells, whereas every synthetic scRNA-seq sample contains the same cells as the single-sample scRNA-seq reference. To make the single-cell variation metrics comparable across samples, each sample is first randomly subsampled to 200 cells. Each cell is downsampled to a common depth by multinomial down-sampling; cells below the target are discarded rather than up-sampled. The target, 161 UMIs, is the 10th percentile of the library sizes of the *real* cells alone, so that it is determined solely from the real multi-sample scRNA-seq and does not change when an approach is added or removed. The conclusions are unchanged at a target depth of 300 UMIs (Supplementary Fig. S3). Samples retaining fewer than 50 cells would be dropped. Because depth is then identical across cells, we use log1p for the normalization.

For the shared embedding used in the sample-mixing analysis, a single principal-component (PC) basis is constructed using cells pooled across all approaches, rather than fitting a separate basis for each approach. To ensure equal contribution across samples in this embedding, each sample is further subsampled to 125 cells, corresponding to the smallest real scRNA-seq sample after rarefaction. From the pooled cells, we select the 300 genes with the highest variance and perform PCA. Because all cells from all approaches are represented in the same PC basis, their coordinates are directly comparable across approaches.

#### Single-cell level: variation metrics

Two complementary metrics are computed: whether cells from different samples are mixed or separated, and how much of a cell’s expression is explained by its sample.

Sample mixing: for each cell we ask whether its neighbourhood in the shared embedding is drawn from one sample or from many, using mLISI (Eq. (12)) over the *S* = 20 sample labels. It is evaluated separately within each approach, so that a cell’s neighbours are only ever cells of the same approach. Values approach 1 when each neighbourhood is drawn from a single sample, so that samples are separable, and 20 when all samples are equally represented, so that samples are interchangeable.

Intraclass correlation: for each gene *g* we fit a one-way random-effects model to its log1p expression across all cells of a group, 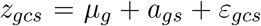, with sample effects *a*_*gs*_ of variance 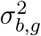 and residuals *ε*_*gcs*_ of variance 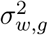, and define

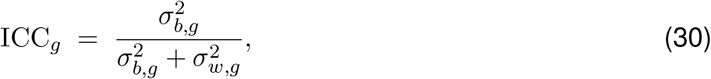

the fraction of that gene’s cell-level variance attributable to sample identity. It is estimated by the analysis-of-variance estimator for unbalanced designs [46],

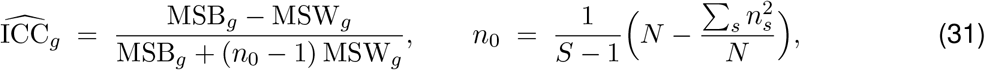

where 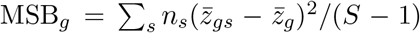 and 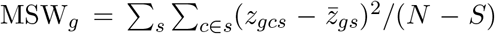 are the between- and within-sample mean squares, *z*_*gcs*_ is the normalized expression of gene *g* in cell *c* of sample 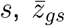 its sample mean, 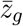 its grand mean, *n*_*s*_ the number of cells in sample *s, S* the number of samples and 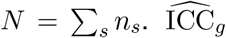 is centered at zero when samples do not differ. Because this ANOVA estimator is not truncated at zero, small negative estimates can occur from sampling variability even though the population ICC is non-negative. We retain these estimates and summarize them by the median across genes, comparing each approach’s median with that of real multi-sample scRNA-seq. Both metrics depend on the number of samples per group, so both use 20 samples per approach, the number of samples in multi-sample scRNA-seq.

### Implementation and software

scStable is implemented as an R package (https://github.com/cfjiang123/scStable) under the MIT license. It depends on `scDesign3` [21] for fitting the scRNA-seq generative model, `Seurat` v5 [31] for preprocessing and clustering, `tmvtnorm` [47] for truncated multivariate Gaussian sampling when **S** is full and `truncnorm` [48] for the diagonal case, `corpcor` [38] for shrinkage covariance estimation, and `nortest` [49] for normality testing in pseudo-count selection.

Default settings follow Steps 1–3, with the pseudo-count *c*_*g*_ chosen by maximizing a normality-test *p*-value (Shapiro–Wilk [50] for *S ≤* 200, Anderson–Darling [51] otherwise; Supplementary Section S1), *λ* = 1, and *R* = 100 synthetic scRNA-seq samples. Per-analysis settings (genes, cells, sample counts, covariance estimator) are summarized in Supplementary Table S1.

Runtime and memory were measured on a standard workstation (Intel Xeon E5-2687W v4 CPU at 3.00 GHz, 251 GB RAM, Ubuntu 22.04.4 LTS). On a single-sample scRNA-seq reference with *G* = 2,000 genes and *C* = 1,614 cells and bulk RNA-seq with *S* = 21 samples, generating *R* = 50 synthetic scRNA-seq samples completed in approximately 19 minutes with a peak memory of *∼*1.6 GB. These resource requirements are compatible with a typical workstation.

## Acknowledgements

We thank Dr. Saidi Wang for initial discussions and explorations on this work.

## Authors’ contributions

C.J. and J.J.L. conceived the project. C.J. implemented the framework and conducted the data analyses. C.J. and J.J.L. interpreted the results, prepared the figures, and drafted and revised the manuscript. J.J.L. supervised the research and manuscript preparation. Both authors read and approved the final manuscript.

## Funding

This work was supported by NIH/NIGMS R35GM140888, NIH/NHGRI R01HG014687, NSF DBI-1846216, and NSF DMS-2113754 to J.J.L.

## Availability of data and materials

The GTEx whole-blood bulk RNA-seq dataset is available through the GTEx Portal (https://gtexportal.org/home). The Zheng4 dataset is distributed through the Bioconductor package DuoClustering2018. The aging mouse lung dataset is available from GEO under accession GSE124872. The gastric cancer dataset is described in Kang et al. [30]; the processed scRNA-seq and matched bulk RNA-seq data used in our analysis were obtained from the original authors’ public GitHub repository (https://github.com/chriscainx/gastric-cancer). Source data un-derlying the figures and supplementary figures, together with the analysis code, are archived on Zenodo at https://doi.org/10.5281/zenodo.23050848. The scStable R package and scripts used to reproduce the analyses and figures are available at https://github.com/cfjiang123/scStable.

## Declarations

### Ethics approval and consent to participate

This study analyzed only previously published, publicly available human and animal transcriptomic datasets. No new human participants were recruited and no new animal experiments were performed. Ethical approvals, informed consent, and animal procedures were reported in the original studies.

### Consent for publication

Not applicable.

### Competing interests

The authors declare that they have no competing interests.

## Notes

### Competing Interest Statement

The authors have declared no competing interest.

