## Supplementary for "scStable: assessing scRNA-seq analysis stability under bulk-derived between-sample variation"

#### S1 scStable model step 2 normalization

For each gene $g$, we choose the pseudo-count $c_g$ in the log transform $x_{gs}^{\log} = \log(x_{gs}^{\mathrm{CPM}} + c_g)$ to make $x_{g1}^{\log}, \ldots, x_{gS}^{\log}$ approximately Gaussian across bulk samples. Specifically, we choose the $c_g$ using a sample-size-dependent test. When $S < 200$ is small, we use the Shapiro–Wilk test; when $S > 200$ is large, we use the Anderson–Darling test. We then select

$$c_g = \arg\max_{c \in \mathcal{C}} p_g(c), \tag{S1}$$

where $p_g(c)$ denotes the $p$-value returned by the chosen normality test applied to $x_{gs}^{\log}$ across samples. Intuitively, this procedure selects the pseudo-count that yields the closest-to-normal log-scale distribution for gene $g$, which supports the Gaussian modeling assumption. In practice, if multiple candidates achieve the same maximal $p$-value, we choose the smallest $c$ to avoid unnecessary distortion of low-expression values.

#### S2 Synthetic sample generation details

We summarize the synthetic scRNA-seq sample generation settings for each analysis in Table S1.

Table S1: Summary of synthetic sample generation settings across analyses.

| **Analysis** | **scRNA-seq** | **Bulk** | **Genes** | **Cells** | **Samples** | **Bulk cov.** |
|---|---|---|---|---|---|---|
| Validation | Zheng4 | GTEx | 200 | All | 100 | Regularized |
| DE: cell type | Zheng4 | GTEx | 2000 | All | 200 | Regularized |
| DE: condition | Lung aging | Same study | 1000 | Macrophage cells | 100 per condition | Diagonal |
| Clustering | Zheng4 | GTEx | 200 | All | 100 per $\lambda$ | Regularized |
| Method comparison | Gastric TME | Paired | 2000 | All | 22 | Diagonal |

Analyses whose bulk RNA-seq reference is GTEx use bulk-reference-free mode; those whose bulk RNA-seq comes from the same study as the scRNA-seq reference (the same individuals for gastric TME, an independent cohort of the same mouse line for lung aging) use bulk-reference mode.

For all analyses, we used the top 10 principal components (PCs) as cell-state covariates for label-free fitting. The GTEx whole-blood dataset ($\sim$800 samples) served as the bulk RNA-seq reference for Zheng4-based analyses, with bulk covariance estimated using a regularized (shrinkage) estimator. For the lung aging and gastric cancer datasets, which have limited study-matched bulk samples, we used a diagonal approximation for the bulk covariance. In the clustering analysis, synthetic samples were generated across a range of sample variation scale factors $\lambda \in [0, 0.05, 0.1, 0.2, 0.3, 0.5, 0.7, 1.0, 1.5, 2.0]$ to enable stress-testing of method stability.

### S3 Between-sample variation comparison

This section supports the comparison of scStable against alternative approaches presented in the corresponding section of the main text, on the paired gastric tumor microenvironment dataset. Supplementary Fig. S1 gives the response of total between-sample variance to the scale factor $\lambda$ across the grid, which is not reproduced in the main figure. Supplementary Figs. S2 and S3 then repeat the pseudo-bulk and the single-cell halves of Fig.3 under a different normalization and a different rarefaction depth, as robustness checks on the two preprocessing choices described in Methods Sections.

Without depth normalization, each pseudo-bulk profile is a plain gene sum over a sample of 202 to 10,658 cells. The leading principal component of the real between-sample deviation matrix then accounts for 87% of the total variance and correlates with log cell number at $r = -0.94$, compared with 59% and $r = -0.04$ after normalization. No approach can reproduce this axis, because every synthetic sample inherits the cell count of the reference sample. The separation between approaches nevertheless holds: in Fig. S2(C) and (D), scStable remains between about 50-fold and five orders of magnitude above ZINB-WaVE, Gaussian noise injection and cell-wise subsampling. The ratios themselves are not interpretable on this scale, because the inflated real denominator lowers every approach: in Fig. S2(D), scStable at $\lambda = 2$ falls from 0.98 to 0.43 of the real total and paired bulk RNA-seq from 0.64 to 0.20, and in Fig. S2(C) paired bulk RNA-seq falls below scStable at $\lambda = 2$ (0.19 against 0.41), whereas it lies above it in Fig.3C (0.80 against 0.61). Likewise, $\rho_c$ in Fig. S2(E) is lower for every approach than in Fig.3E, but scStable remains well separated from the alternatives: it reaches $\rho_c = 0.807$ at $\lambda = 2$ and 0.564 at $\lambda = 1$, against 0.597 for paired bulk RNA-seq and 0.145, 0.107 and 0.030 for ZINB-WaVE, cell-wise subsampling and Gaussian noise injection.

Both metrics in Fig. S3 behave as they do at 161 UMIs. Real multi-sample scRNA-seq reaches an mLISI of 4.00 and scStable 4.23 at $\lambda = 1$ and 2.58 at $\lambda = 2$, whereas ZINB-WaVE, Gaussian noise injection and cell-wise subsampling reach 8.64, 8.76 and 8.78. In the median gene, 4.3% of the cell-level variance is attributable to sample identity in real multi-sample scRNA-seq and 3.3% and 6.9% under scStable at $\lambda = 1$ and $\lambda = 2$, whereas for the three alternatives it is indistinguishable from zero. The single-cell conclusions therefore do not depend on the rarefaction depth.

### S4 Stability-based hyperparameter selection for Louvain clustering

To determine whether scStable can support data-specific tuning of clustering hyperparameters, we evaluated Louvain clustering across a grid of combinations of the number of principal components ($n_{\mathrm{PC}}$), the number of nearest neighbors ($k_{\mathrm{NN}}$), and the resolution parameter ($r$). For each combination, Louvain clustering was applied to the single-sample scRNA-seq reference and to each scStable synthetic sample generated under realistic between-sample variation ($\lambda = 1$). Stability was assessed by comparing the clustering labels obtained from the synthetic samples with those obtained from the reference sample under the same hyperparameter setting, using the four agreement metrics described in the main text.

Supplementary Fig. S4 shows that clustering stability varies substantially across hyperparameter combinations. All four metrics identify the same stable setting. For retrospective evaluation only, we additionally compare the reference clustering obtained under each hyperparameter combination with the known cell-type labels. The stable setting selected by scStable also shows high agreement with the true labels, supporting the use of stability under bulk-derived between-sample variation for label-free hyperparameter selection.

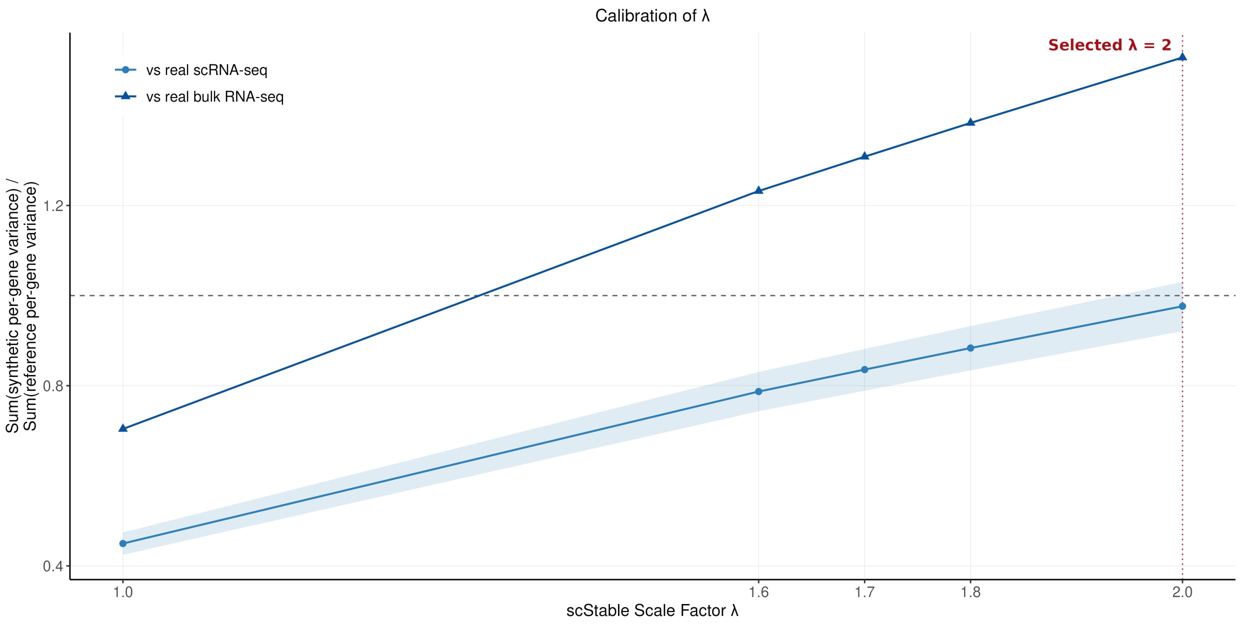


Figure S1: **Sensitivity of between-sample variance to the scStable scale factor $\lambda$.** Total between-sample variance summed over the 1,369-gene panel, carried by scStable synthetic samples relative to real multi-sample scRNA-seq (circles) and to real multi-sample bulk RNA-seq (triangles), as a function of $\lambda$. The dashed line marks exact agreement (ratio $= 1$), the shaded band is the 95% gene-bootstrap interval for the scRNA-seq analysis, and the dotted line marks $\lambda = 2$, the setting at which the scRNA-seq between-sample variance ratio is closest to one. Against real multi-sample scRNA-seq the ratio rises monotonically from 0.450 at $\lambda = 1$ to 0.976 at $\lambda = 2$. Against real bulk RNA-seq it rises from 0.704 to 1.528 and crosses one already at $\lambda \approx 1.3$. No single $\lambda$ matches both references, and $\lambda = 1$ and $\lambda = 2$ are both reported in the main text as a sensitivity analysis.

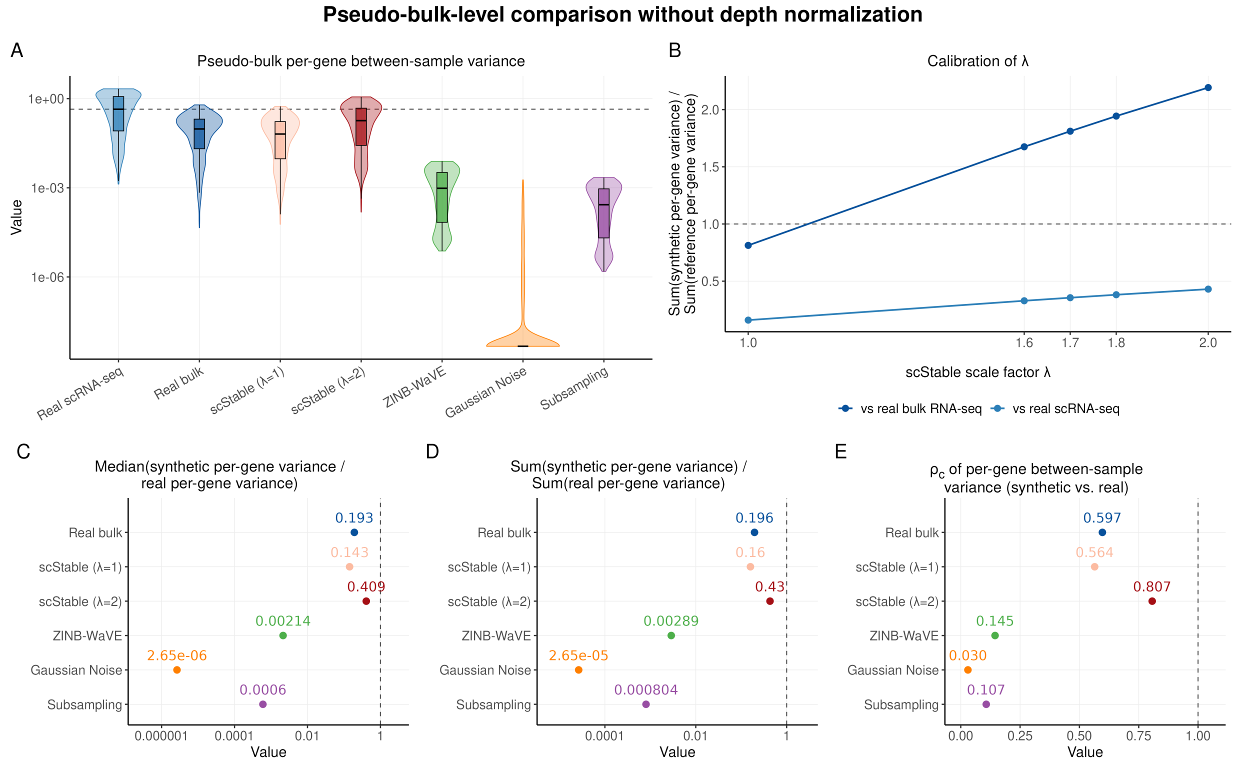


Figure S2: **Pseudo-bulk comparison of between-sample variation without depth normalization.** Panels (A) and (C)–(E) repeat Fig.3 B–E on raw pseudo-bulk sums, without equalizing library size across samples; (B) is the counterpart of Supplementary Fig. S1 on the same scale. **(A)** Distributions of per-gene between-sample variance for real multi-sample scRNA-seq, paired bulk RNA-seq, and synthetic scRNA-seq samples generated by each approach. The dashed line indicates the median for real multi-sample scRNA-seq. **(B)** Total between-sample variance of scStable synthetic scRNA-seq samples relative to real multi-sample scRNA-seq and to paired bulk RNA-seq, as a function of the scale factor $\lambda$. The dashed line at 1 indicates exact agreement. **(C)** Median across genes of the ratio of each approach's per-gene between-sample variance to that of real multi-sample scRNA-seq. **(D)** Total between-sample variance relative to real multi-sample scRNA-seq. In (C,D), the dashed line at 1 indicates exact agreement. **(E)** $\rho_c$ between $\log_{10}$ per-gene between-sample variances from each approach and those from real multi-sample scRNA-seq; the dashed line at 1 indicates exact agreement.

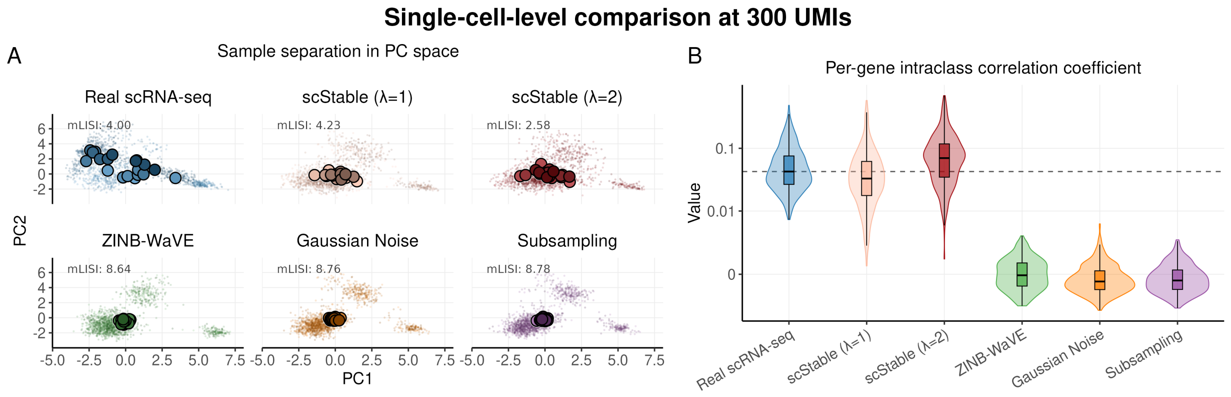


Figure S3: **Single-cell-level between-sample variation at a deeper rarefaction target (**300 **UMIs per cell).** Panels (A) and (B) repeat Fig.3 F and G after rarefying every cell to 300 instead of 161 UMIs. **(A)** Cells from each approach projected into a common principal-component (PC) space fitted on their pooled cells. Small points represent individual cells and large outlined points represent sample centroids in the PC space. For comparability, 20 samples are used for each approach. mLISI is calculated over sample identity and ranges from 1, when samples are locally separated, to 20, when samples are locally indistinguishable. **(B)** Per-gene intraclass correlation coefficients (Eq. 30), measuring the fraction of cell-level expression variance attributable to sample identity. The dashed line indicates the median for real multi-sample scRNA-seq.

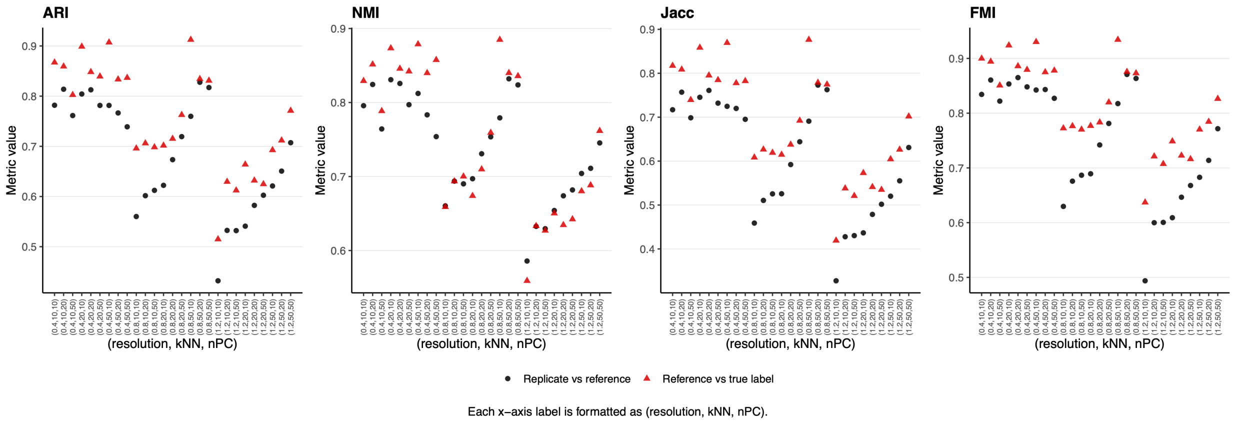


Figure S4: **Stability-based selection of Louvain clustering hyperparameters.** Louvain clustering was evaluated across a grid of hyperparameter combinations spanning $n_{\text{PC}} \in \{10, 20, 50\}$, $k_{\text{NN}} \in \{10, 20, 50\}$, and $r \in \{0.40, 0.80, 1.20\}$ using scStable synthetic samples generated at $\lambda = 1$. Each panel shows one of four clustering agreement metrics: Adjusted Rand Index (ARI), Normalized Mutual Information (NMI), Jaccard Index (Jacc), and Fowlkes–Mallows Index (FMI). Hyperparameter combinations are arranged along the $x$-axis and formatted as $(r, k_{\text{NN}}, n_{\text{PC}})$. Black points indicate stability, quantified as the agreement between clustering labels from the synthetic samples and those obtained from the single-sample scRNA-seq reference under the same hyperparameter setting. Red triangles indicate agreement between the reference clustering and the known cell-type labels and are shown only for retrospective evaluation. We could select the stable hyperparameter combination based on the highest stability across metrics without using the cell-type labels.
